# Mechanisms underlying proximity between oral commensal bacteria

**DOI:** 10.1101/2020.09.29.318816

**Authors:** Dasith Perera, Anthony McLean, Viviana Morillo Lopez, Kaileigh Cloutier-Leblanc, Eric Almeida, Kiana Cabana, Jessica Mark Welch, Matthew Ramsey

## Abstract

Complex polymicrobial biofilm communities are abundant in nature particularly in the human oral cavity where their composition and fitness can affect health. While the study of these communities during disease is essential and prevalent, little is known about interactions within the healthy plaque community. Here we describe interactions between two of the most abundant species in this healthy microbiome, *Haemophilus parainfluenzae* and *Streptococcus mitis*. We discovered that *H. parainfluenzae* typically exists adjacent to Mitis group streptococci *in vivo* with which it also positively correlated based on microbiome data. By comparing *in vitro* coculture data to *ex vivo* microscopy we revealed that this co-occurrence is density dependent and further influenced by H_2_O_2_ production. We discovered that *H. parainfluenzae* has a more redundant, multifactorial response to H_2_O_2_ than related organisms and that the integrity of this system enhances streptococcal fitness. We also show that Mitis group streptococci can act as an *in vivo* source of NAD for *H. parainfluenzae* and that streptococci *in vitro* evoke patterns of carbon utilization from *H. parainfluenzae* that are similar to those observed *in vivo*. Our findings describe mechanistic interactions between two of the most abundant and prevalent members of healthy supragingival plaque that contribute to their survival *in vivo*.

## Introduction

Within the human oral microbiome, supragingival plaque (SUPP) is a polymicrobial biofilm that grows on the tooth surface above the gum line. The composition of SUPP has been long studied beginning with Antony von Leeuwenhoek in 1683 (Mikx 1983) and resolved in great detail both in composition by microbiome studies (Dewhirst et al. 2010; Human Microbiome Project Consortium 2012; Eren et al. 2014) and physical structure by microscopy and attachment-based studies (Mark Welch et al. 2016; Kolenbrander et al. 2002). *Haemophilus parainfluenzae* is one of the most abundant and prevalent species in the SUPP of healthy individuals (Human Microbiome Project Consortium 2012; Eren et al. 2014). Likewise *Streptococcus* is one of the most abundant and prevalent genera in this environment with species within the Mitis group (*S. mitis, S. oralis, S. australis, S. infantis*, and others) (Zheng et al. 2016; Jensen, Scholz, and Kilian 2016) being particularly abundant (Liljemark et al. 1984; Dewhirst et al. 2010; Eren et al. 2014). In addition to SUPP, these organisms are highly abundant in other oral and extraoral sites. Despite these being highly abundant and prevalent species in the oral microbiome, it is unknown if they exist in close enough contact to influence each other and if so, by what mechanism(s).

In this study we demonstrate that in SUPP *H. parainfluenzae* grows in close proximity to Mitis group streptococci. Many oral *Streptococcus spp*. are known to produce anti-microbial substances, including hydrogen peroxide (H_2_O_2_) in aerobic conditions. Therefore, any bacterium adjacent to these *Streptococcus sp*. aerobically would need to have adapted the ability to withstand H_2_O_2_ (L. Zhu and Kreth 2012; Ramsey, Rumbaugh, and Whiteley 2011). Thus, H_2_O_2_ production by *Streptococcus spp*. can be an important mechanism driving community composition and spatial arrangement. H_2_O_2_ produced by *S. pneumoniae* has been demonstrated to inhibit growth of the respiratory tract pathogens *Moraxella catarrhalis, Neisseria meningitidis* and *H. influenzae* (Pericone et al. 2000), and mechanisms of responding to H_2_O_2_ have been established for many species including *H. influenzae* (Harrison, Bakaletz, and Munson 2012) but not *H. parainfluenzae*. Likewise, most *Streptococcus sp*. also produce lactic acid as a metabolic end product of carbohydrate fermentation (Kreth, Merritt, and Qi 2009) which can support the growth of some species (van der Hoeven, Toorop, and Mikx 1978; Mikx and Van der Hoeven 1975) while excluding others (Mashimo et al. 1985). The ability of *Streptococcus sp*. to rapidly consume high-energy carbohydrates while producing lactic acid and H_2_O_2_ provides it with a competitive advantage and it is currently unknown how *H. parainfluenzae* tolerates these stresses or how this relates to its existence *in vivo*.

Here we demonstrate that predicted associations between *H. parainfluenzae* and Mitis group streptococci occur *in vivo* as reflected in microscopy analysis of *ex vivo* samples. We further show that *S. mitis* can kill *H. parainfluenzae* by H_2_O_2_ production in a dose-dependent manner which is reflected *in vivo* with an apparent density-dependent association between the two taxa. We also observed that the sole catalase gene product of *H. parainfluenzae* plays only a minor role in H_2_O_2_ resistance, in contrast to other catalase-positive species including *H. influenzae* (Juneau et al. 2015; Bishai et al. 1994). We then assessed the contribution of several gene products that provide H_2_O_2_ resistance revealing that they too provide only modest levels of protection, suggesting a redundant, multifactorial mechanism in H_2_O_2_ resistance. Despite this antagonism, *H. parainfluenzae* repeatedly was found directly adjacent to *Streptococcus sp. in vivo* which is supported by our finding that Mitis group streptococci are substantial producers of NAD which *H. parainfluenzae* cannot synthesize on its own nor obtain from host saliva. Comparisons of *in vitro* interaction data with *in vivo* metatranscriptome studies reveal *H. parainfluenzae* changes in carbon source utilization and other behaviors indicating that these are likely due to interactions with Mitis group members *in vivo*. These results provide a robust characterization of *H. parainfluenzae’s* role in the oral microbiota and reveal ways it has evolved to exist alongside streptococci in the oral cavity and likely beyond. This study details interactions between two prominent members of a complex natural biofilm community and allow us to demonstrate highly detailed mechanisms of interaction that help drive micron-scale arrangements between these organisms that is likely conserved in other host sites where they overlap.

## Results

### *H. parainfluenzae* co-occurs with *S. mitis* and related streptococci in human supragingival plaque

*H. parainfluenzae* is one of the most abundant species in healthy supragingival plaque (Eren et al. 2015). To study multispecies interactions of *H. parainfluenzae* with healthy oral commensals, we needed to determine which species it was most likely to interact with. To examine this, we used species-specific microbiome data from a 117 subjects sampled by the Human Microbiome Project (HMP) to predict species-species interactors with *H. parainfluenzae* (Fig. 1). A re-analysis of HMP species-assigned metagenomic data indicated that *H. parainfluenzae* is an abundant and prevalent member of SUPP detected in all 117 subjects averaging 7.6% relative abundance based on sequence reads (Fig. 1). We compared the upper quartile (n=29) of these subjects ranked by highest *H. parainfluenzae* abundance to the remainder of subjects (n=88) via LEfSe analysis (Segata et al. 2011) to determine which species were significantly likely to co-occur with *H. parainfluenzae* (Fig. 1). Interestingly, this indicated that individuals enriched in *H. parainfluenzae* also have an abundance of *Streptococcus* sp. especially those belonging to the Mitis group. This includes the species *S. australis*, *S. infantis*, *S. pneumoniae*, *S. oralis*, *S. peroris*, and *S. mitis*. We therefore decided to investigate *H. parainfluenzae* interactions with *S. mitis* as a representative of the Mitis group in order to examine the mechanisms of taxon-taxon interactions.

**Figure 1:**
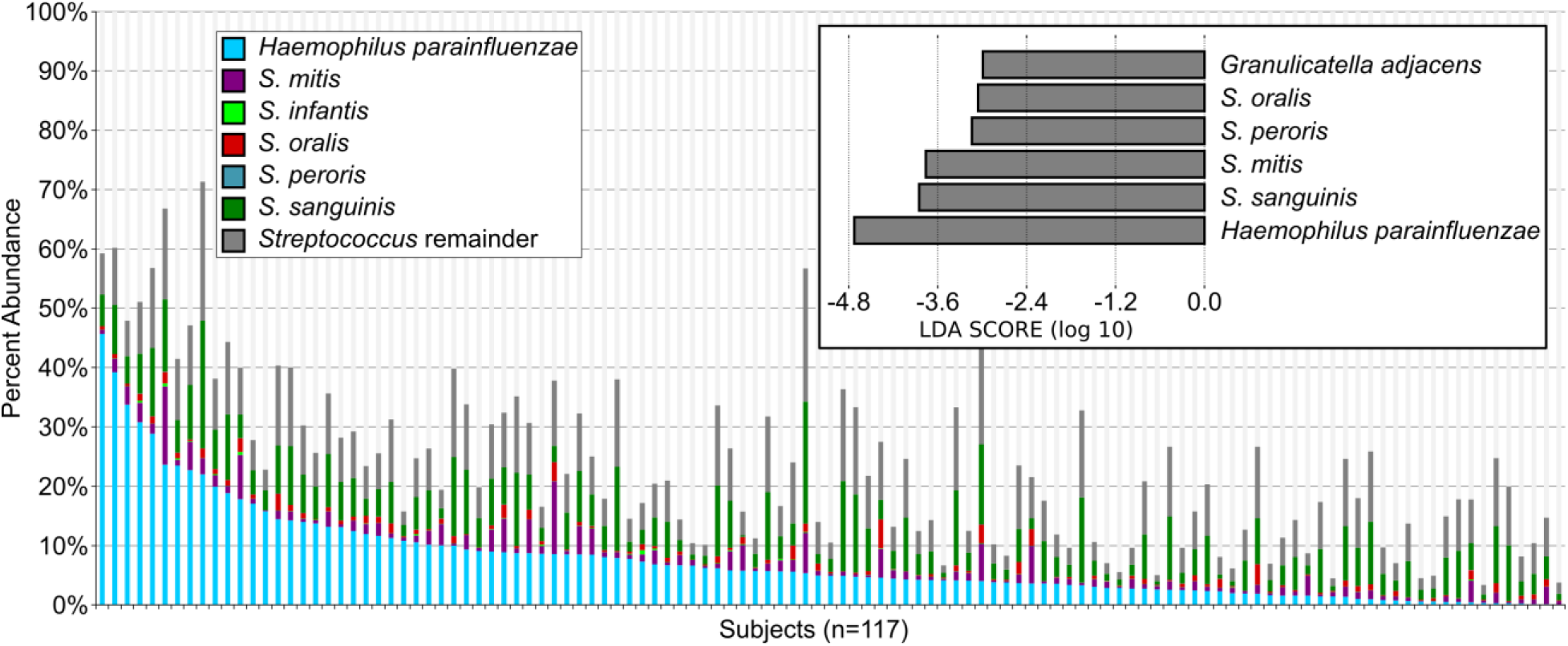
Read abundance data and predicted correlations between taxa in supragingival plaque. Human Microbiome Project (HMP) metagenome data of supragingival plaque was used to plot the relative abundance and prevalence of species of interest including *Haemophilus parainfluenzae* (red), several Mitis group streptococci and all remaining *Streptococcus* spp. (dark grey). (Top Right) The top 25% of subjects based on *H. parainfluenzae* abundance were compared to the remainder via LEfSe analysis. Shown are species enriched in this comparison above an LDA score ≤ −3.0.

### Species-specific FISH demonstrates frequent *H. parainfluenzae* - *S. mitis* co-proximity

Within the supragingival plaque, *S. mitis* and related streptococci are a frequent feature of *H. parainfluenzae’s* micron-scale environment. Visualizing these species with FISH probes showed that most *H. parainfluenzae* in supragingival plaque are located within a few micrometers of cells labeled with a probe that hybridizes with *S. mitis, S. oralis*, and *S. infantis* (Fig. 2, Table 3, hereafter referred to as “*S. mitis*”). The median distance separating a *H. parainfluenzae* cell from the nearest *S. mitis* cell is 1.14 μm, and 92% of *H. parainfluenzae* cells in the plaque images fall within 10 μm of the nearest *S. mitis* cell (Fig. 2A). Due to the proximity of these taxa, H_2_O_2_ and other inhibitory or promotive compounds excreted by *S. mitis* could reasonably be expected to perfuse the substrate in which most *H. parainfluenzae* grow.

**Figure 2.**
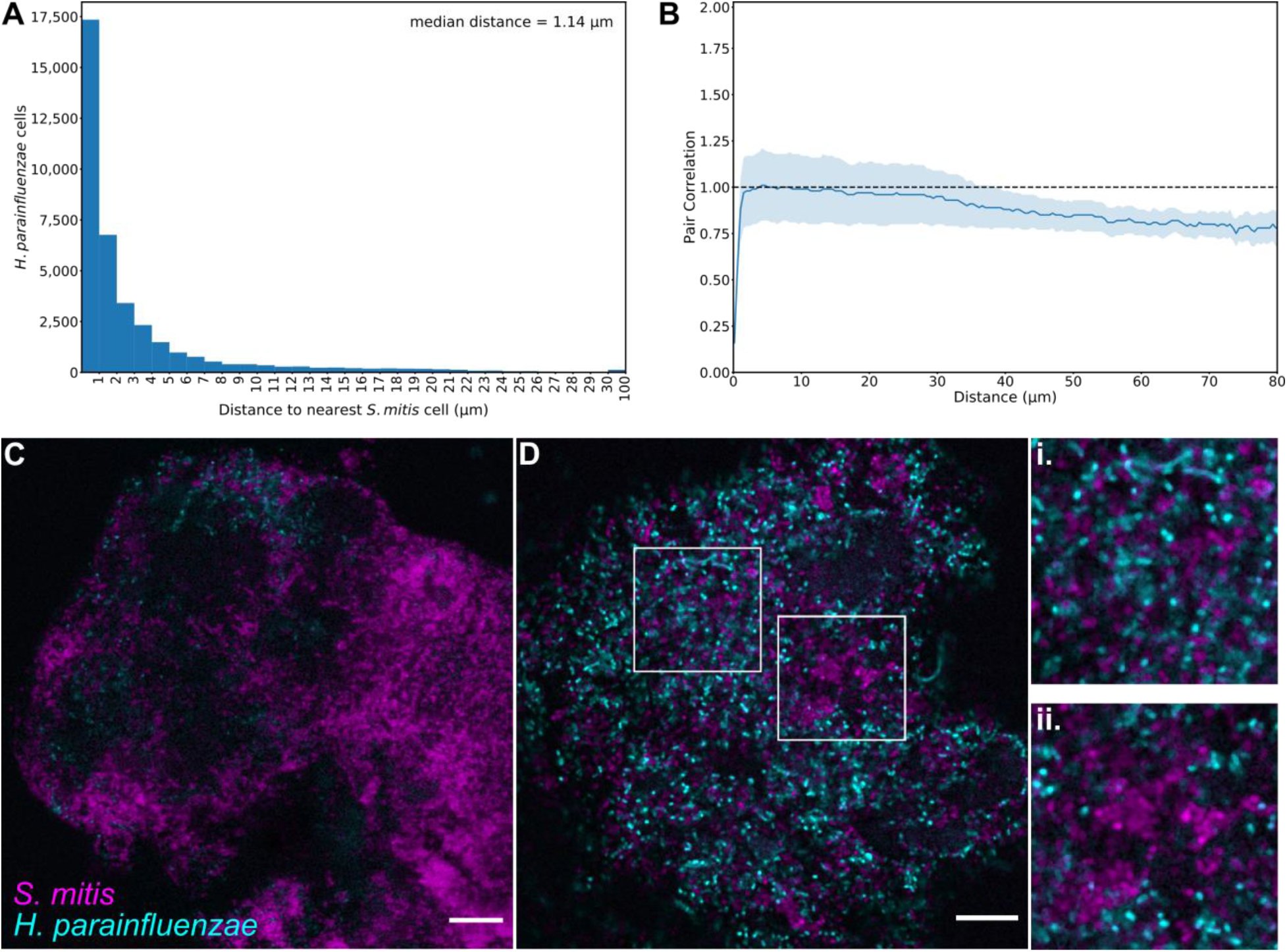
*H. parainfluenzae* distribution is related to the density of *S. mitis in vivo*. (A) Histogram of the distance to the nearest *S. mitis* cell, measured edge-to-edge, from each of the 37,591 *H. parainfluenzae* cells in 41 fields of view. (B) Pair correlations between *H. parainfluenzae* and *S. mitis* cells. The lighter lines represent the bounds of the 95% confidence interval for the correlation values. The dashed line represents the null hypothesis: the pair correlation equals one. n = 41 fields of view. (C) Plaque with sparsely distributed *H. parainfluenzae* (cyan). (D) Plaque with high abundances of both *S. mitis* (magenta) and *H. parainfluenzae*. (i) Most *H. parainfluenzae* cells are within a few microns of the nearest *S. mitis* cell. Generally, *H. parainfluenzae* are randomly distributed with respect to *S. mitis*. (ii) *H. parainfluenzae* avoids the highest densities of *S. mitis*. Scale bars indicate 10 μm.

The effect of each species on the growth of the other *in vivo* is likely to be reflected in the micron-scale spatial organization of the taxa relative to one another. We used image analysis with the method of linear dipoles (Daims, Lücker, and Wagner 2006; Daims and Wagner 2011) to evaluate the spatial cross-correlation between the taxa. For dipole lengths ranging from 1 to 39 μm, the pair correlations did not significantly differ from the case of random distribution (Fig. S1). This result indicates that when *H. parainfluenzae* is present within a region of plaque, it is randomly distributed with respect to *S. mitis*. For both the very shortest (0.15 to 0.6 μm) and longest (39.3 to 90.15 μm) dipole lengths, the two species had a significantly negative correlation (p < 0.05). The former would be expected due to the effect of spatial exclusion as multiple cells cannot occupy the same space in a single focal plane. The latter is consistent with the observed occurrence of regions of plaque without either taxon.

In contrast to the apparently random distribution of *H. parainfluenzae* with respect to *S. mitis* across entire images, areas with the highest densities of *S. mitis* showed a reduced density of *H. parainfluenzae*. To assess the spatial organization of the two taxa in regions of high *S. mitis* density, we divided images into 6.64 μm by 6.64 μm blocks and compared the local densities of both taxa at this resolution. The mean density of *H. parainfluenzae* increases as the percent of a block covered by *S. mitis* increased from 0 to around 20% (Fig. S1). This trend is likely due, at least in part, to the effect of variation within the overall quantity of plaque in each block because the density of each taxon has a positive linear relationship with the overall plaque density. As the density of *S. mitis* increases above 20%, however, the mean *H. parainfluenzae* density decreases. While the mean *H. parainfluenzae* densities for blocks with *S. mitis* densities greater than 25% have a high degree of error, a qualitative assessment of the plaque images with the highest density clumps of *S. mitis* supports the conclusion (Fig. 2C,D). The finding that the mean density of *H. parainfluenzae* begins to decline once the *S. mitis* densities exceeded a threshold value suggests a *S. mitis*-density-dependent inhibitory effect on *H. parainfluenzae* growth. Given that both microbiome sequencing and microscopy imaging of *in vivo* supragingival plaque samples indicate significant co-occurrence and co-proximity between *H. parainfluenzae* and *S. mitis* (Figs. 1,2), it is important to determine the mechanisms that dictate these observations.

### *S. mitis* eliminates *H. parainfluenzae* via production of H_2_O_2_

We used a reductionist approach to investigate the growth of each organism *in vitro* in close proximity via a colony biofilm model (Merritt, Kadouri, and O’Toole 2005; Ramsey et al. 2016). We inoculated *H. parainfluenzae* and *S. mitis* in mono and coculture on a BHIYE-HP agar plate for 24 hours and then measured growth yields by CFU counts. Cocultures inoculated at equal densities revealed that *S. mitis* strongly reduced *H. parainfluenzae* numbers (Fig. 3) nearly 100-fold below inoculum density, indicating active killing. This effect is dose-dependent as we observed when inoculums of *S. mitis* were either equivalent or 3-fold greater than *H. parainfluenzae* there was a significant reduction in the growth yield of *H. parainfluenzae* compared to monoculture. However, when *S. mitis* inoculum was 10-fold lower than *H. parainfluenzae*, there was no significant change in growth yield compared to monoculture

**Figure 3:**
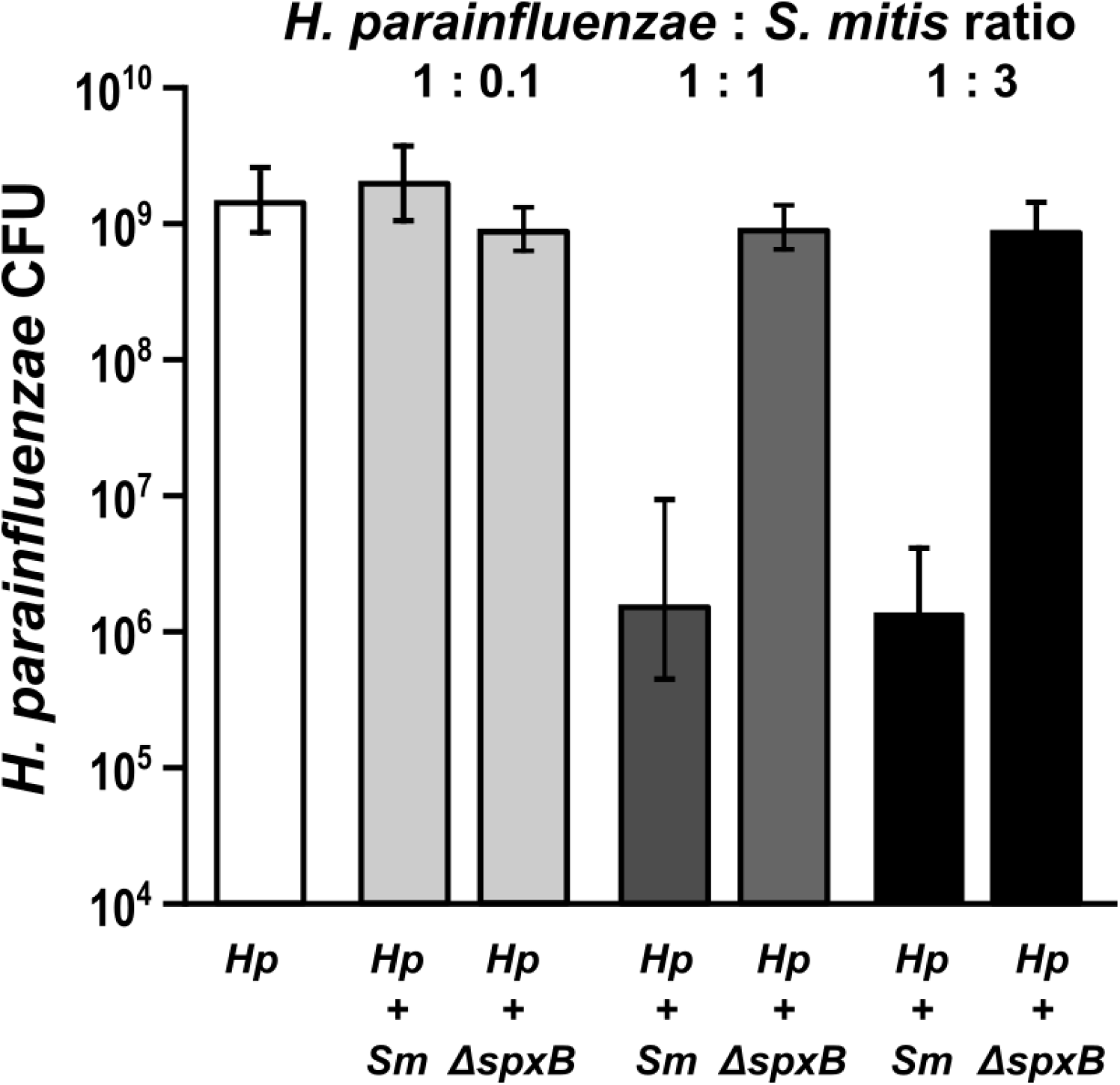
*H. parainfluenzae* growth is inhibited by *Streptococcus mitis* produced H_2_O_2_ in a dose dependent manner. *H. parainfluenzae (Hp*) CFU counts in mono and coculture with wildtype (*Sm*) or a pyruvate oxidase mutant (*ΔspxB*) of *S. mitis. Hp* had an initial inoculum using 10μL at an OD_600_ of 1, which corresponds to 4.65×10^6^ CFU/ml. Wildtype (*Sm*) and *S. mitis ΔspxB* with initial inoculums using 10μL at an OD_600_ of either 0.1, 1 or 3 which corresponds to an average of 2.45×10^5^, 1.55×10^6^ or 3.45×10^6^ CFU/ml. Data are mean CFU counts with error bars indicating standard deviation for n≥3. *denotes p< 0.001 using a Student’s t-test compared to monoculture.

This dose-dependent reduction was abolished when *H. parainfluenzae* was cocultured with a strain of *S. mitis* lacking pyruvate oxidase (*ΔspxB*) and unable to produce H_2_O_2_. Figure 3 demonstrates that *H. parainfluenzae* growth yield when cocultured with *ΔspxB* is not significantly different from monoculture at any ratio tested, indicating that *S. mitis-*produced H_2_O_2_ is responsible for *H. parainfluenzae* inhibition. Additionally, *H. parainfluenzae* was unable to grow in supernatants of *S. mitis* culture unless pre-treated with exogenous catalase (data not shown), further supporting the finding that H_2_O_2_ production limits *H. parainfluenzae* growth in these conditions. Therefore, while our *ex vivo* data shows co-occurrence/co-proximity between these taxa and suggests a *S. mitis*-density-dependent association, our *in vitro* coculture data suggests that H_2_O_2_ toxicity can limit these interactions.

### Individual H_2_O_2_ sensitive genes do not affect the fitness of *H. parainfluenzae* in coculture with *S. mitis*

Given that our *ex vivo* data suggests that *H. parainfluenzae* has co-evolved in proximity to H_2_O_2_-producing Streptococci, we decided to investigate the involvement of known H_2_O_2_ relevant gene products present on the *H. parainfluenzae* genome by constructing gene deletions and assessing fitness after H_2_O_2_ exposure. *H. parainfluenzae* possesses an *oxyR* coding sequence whose gene product is the global transcriptional regulator that responds to H_2_O_2_ in many species. Also present is a single catalase gene (*katA*) shown to be essential for H_2_O_2_ resistance in *H. influenzae* (Bishai et al. 1994). Additionally, *H. parainfluenzae* possesses a cytochrome-C peroxidase (*ccp*), crucial for peroxide resistance in other *Pasteurellaceae* (Takashima and Konishi 2008), *Campylobacter jejuni* (Ishikawa et al. 2003), as well as *H. influenzae* (Wong et al. 2007). We also made deletions of the peroxiredoxin (*prx*), peroxiredoxin-glutaredoxin (*pdgX*) and glucose-6-phosphate dehydrogenase (*g6p*) genes that are known to contribute to oxidative stress protection in *H. influenzae* and other species (Juneau et al. 2015; Izawa et al. 1998; Lundberg et al. 1999; Perkins et al. 2015).

We quantified changes in H_2_O_2_ resistance for each mutant using a zone of inhibition assay (Fig. 4A). The largest increase in sensitivity as expected was observed following deletion of *oxyR*, while all other individual gene deletion mutants demonstrated more modest sensitivities to H_2_O_2_. Double mutants for *katA* and *ccp* demonstrated more significant increases in sensitivity vs individual gene mutants indicating a combinatorial effect of H_2_O_2_ protective gene products. Surprisingly, MIC concentrations for many of these genes were nearly identical (Table S1) indicating that the individual contributions of these gene products to H_2_O_2_ tolerance are too minimal for MIC assay resolution. We next tested the fitness of each mutant in coculture with H_2_O_2_-producing *S. mitis* (Fig. 4B). While OxyR was shown to be essential for *H. parainfluenzae* survival in coculture, deletion of individual genes typically controlled by OxyR showed no significant difference when compared to wildtype which contrasts greatly to similar mutants in *H. influenzae* (*katA*) (Bishai et al. 1994) and other species. However, the *ΔkatA+Δccp* mutants’ growth yield was significantly inhibited in *S. mitis* coculture indicating an additive effect of these gene products on H_2_O_2_ resistance. These data demonstrate that the mechanism of *H. parainfluenzae* resistance to H_2_O_2_ involves a complex multifactorial system unlike *H. influenzae* that is under further investigation by our group.

**Figure 4:**
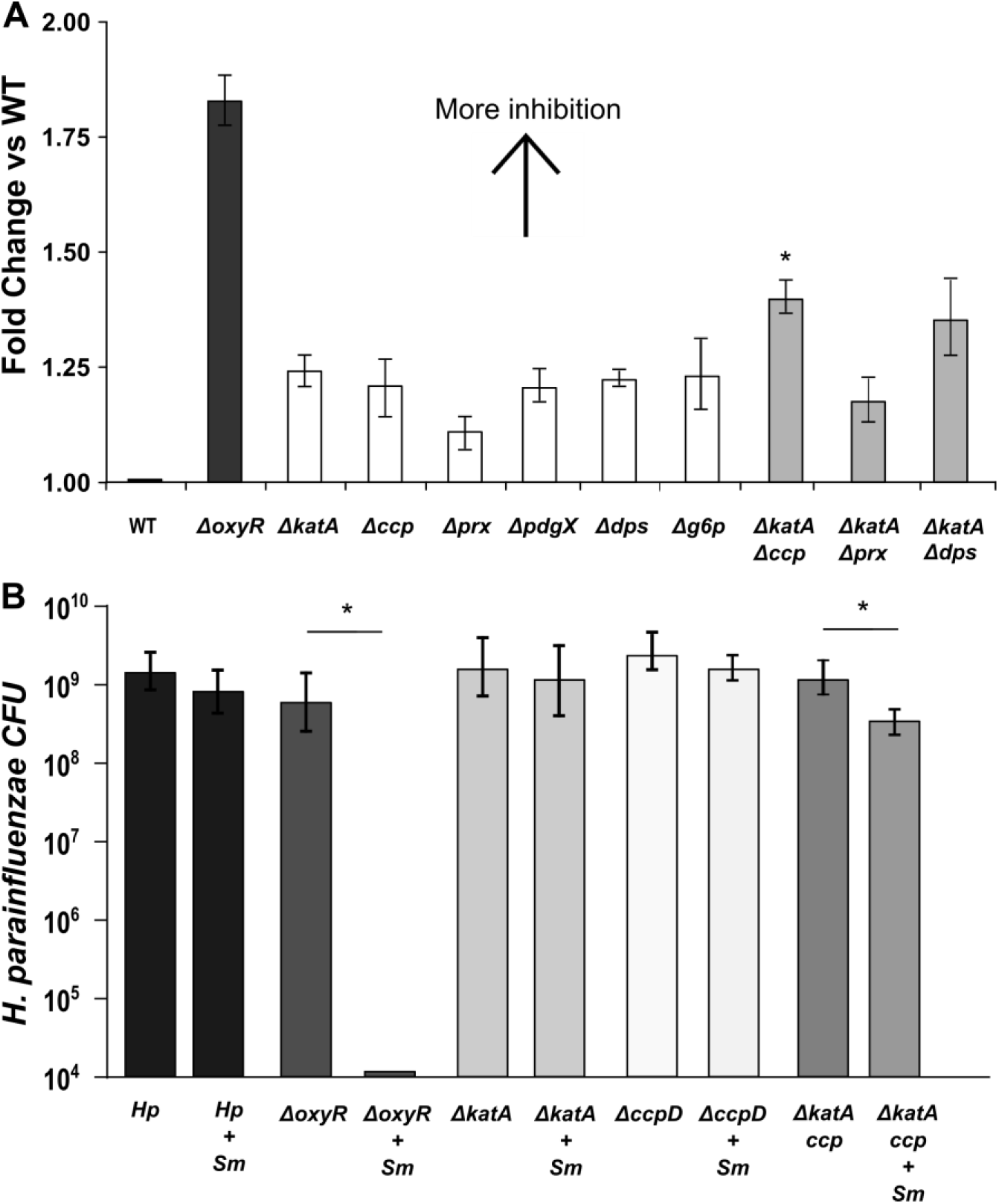
*H. parainfluenzae* resistance to H_2_O_2_ relies on the contribution of multiple genes. We assessed the sensitivity of wildtype *H. parainfluenzae* (WT) to H_2_O_2_ or coculture yield with *S. mitis* vs mutants lacking *oxyR*, catalase (*katA*), cytochrome c peroxidase (*ccp*), peroxiredoxin (*prx*), glutaredoxin-peroxiredoxin (*pdgX*), DNA-binding protein from starved cells (*dps*) and glucose-6-phosphate dehydrogenase (*g6p*). (A) We spotted 5μL of 30% H_2_O_2_ on a paper disk on an agar plate lawn of *H. parainfluenzae*. Zones of inhibition (areas of no visible *H. parainfluenzae* growth) were measured after 24 hours and fold change calculated compared to wildtype. Data are the mean fold change relative to WT; error bars indicate standard deviation for n≥3. All strains were significantly different from WT, while *ΔkatA-Δccp* was significantly different from *ΔkatA* (padj < 0.05), based on t-test with Bonferroni correction. (B) WT and mutant *H. parainfluenzae* CFU following 24h coculture with *S. mitis*. *H. parainfluenzae* inoculums were 4.65 ×10^6^ CFU/ml. *S. mitis* inoculum was 2.45×10^5^ CFU/ml. Data are represented as mean CFU, error bars indicate standard deviation for n≥3. *denotes p< 0.05 by Student’s t-test compared to monoculture.

### H_2_O_2_ detoxification by *H. parainfluenzae* supports *S. mitis* growth

Cocultures revealed that *H. parainfluenzae* increased the growth yield of *S. mitis* 162-fold when at equal ratios (Fig. S2A). Because streptococci can limit their own growth due to H_2_O_2_ accumulation (Xu, Itzek, and Kreth 2014), we hypothesized that *H. parainfluenzae* increases the growth of *S. mitis* via H_2_O_2_ detoxification. To test this hypothesis we compared mono and coculture yields following the addition of (20 U/ml) exogenous catalase (Fig. S2B). We observed that the 162-fold growth benefit for *S. mitis* was decreased to only 16-fold when catalase was added indicating that *H. parainfluenzae* H_2_O_2_ detoxification accounts for the majority of *S. mitis’* coculture benefit. This is further demonstrated by the inability of *H. parainfluenzae ΔoxyR* to aid *S. mitis* growth (Fig. S2A). Using the *S. mitis ΔspxB* strain which is unable to produce H_2_O_2_ we also observed a more modest growth benefit in coculture compared to its wild-type (Fig. S2C). Just as *S. mitis* density affected *H. parainfluenzae* growth yield (Fig. 3) we also observed that *H. parainfluenzae* density also affected *S. mitis* growth yield. When *S. mitis* / *H. parainfluenzae* ratios were 1:1 or less, *S. mitis* growth yield was greater vs monoculture (Fig. S2C). These data indicate that H_2_O_2_ detoxification by *H. parainfluenzae* provides the majority of *S. mitis* growth enhancement so long as *S. mitis* density is not too high.

### *S. mitis* and other *Streptococcus sp*. support *H. parainfluenzae* growth

Like other *Haemophilus sp., H. parainfluenzae* is a NAD auxotroph and must exist in environments where NAD, nicotinamide mononucleotide (NMN) or nicotinamide riboside (NR) are supplied by the host or other microorganisms (Cynamon, Sorg, and Patapow 1988). We determined that sterile human saliva supplemented with glucose was unable to support *H. parainfluenzae* growth unless NAD was added (data not shown) suggesting that adjacent microbes are an important source of NAD for *H. parainfluenzae* in the oral cavity. As *Corynebacterium* and *Streptococcus* are two of the most abundant genera in tooth plaque we tested the ability of species from both genera to complement NAD auxotrophy. While supernatants of all species tested supported modest growth (data not shown), spot assays using actively growing cells in close proximity to *H. parainfluenzae* lawns on agar plates lacking NAD showed robust growth of *H. parainfluenzae* adjacent to *S. mitis* and *S. sanguinis* but no other taxa (Fig. 5). These data suggest that *H. parainfluenzae* can obtain NAD specifically from these two taxa when they are in close proximity. It is interesting that the two species that significantly correlate with *H. parainfluenzae* in microbiome data (Fig. 1) enhance its growth most strongly in the absence of NAD.

**Figure 5:**
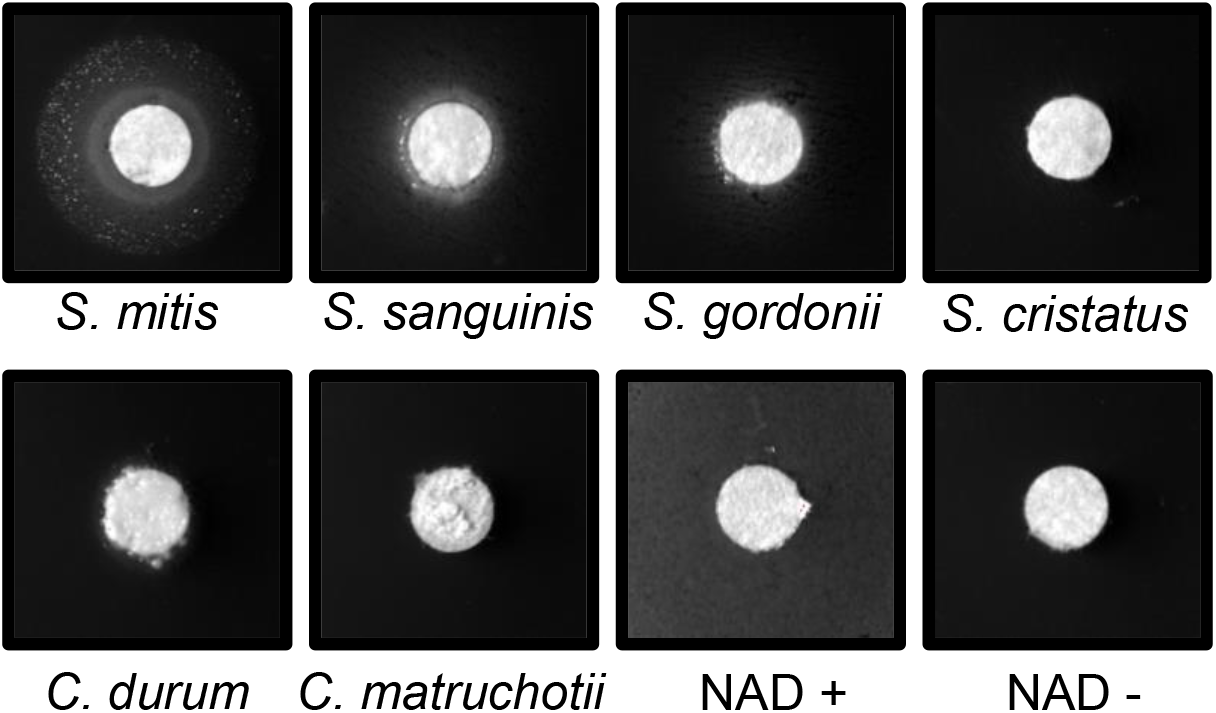
*Streptococcus-produced* Nicotinamide Adenine Dinucleotide (NAD) supports *H. parainfluenzae* growth. Cultures of *S. mitis, S. sanguinis, S. gordonii, S. cristatus, Corynebacterium matruchotii* and *C. durum* were grown overnight, normalized based on optical density, and spotted onto paper discs over lawns of *H. parainfluenzae* spread on solid agar medium lacking NAD. Plates were incubated for 48h before growth was observed. Lighter rings close to the disc indicate *H. parainfluenzae* growth. NAD+ indicates addition of 11 mM NAD as a positive control.

### *In vitro* transcriptional responses of *H. parainfluenzae* to *S. mitis*

To further examine the mechanisms of interaction that take place between these two species, we investigated the transcriptional responses of *H. parainfluenzae* when cocultured with *S. mitis* aerobically. *S. mitis* transcriptional responses to *H. parainfluenzae* are the focus of a separate study. We observed 387 *H. parainfluenzae* significantly differentially expressed genes greater than 2-fold in coculture (Appendix 1), compared to monoculture. Interestingly, we did not observe an increase in *H. parainfluenzae* catalase in coculture; however, based on transcript abundance catalase appeared to be well expressed in mono and coculture conditions. Among genes that are typically involved in oxidative stress responses, *dps* had a 2.2-fold increase in coculture, suggesting that *H. parainfluenzae* is sequestering free intracellular Fe^2+^, potentially to prevent oxidative damage. Surprisingly however, a number of other genes involved in H_2_O_2_ were repressed in coculture, including *ccp, pdgX*, thioredoxin (*trxA*), glutaredoxin (*grx*) and thiol peroxidase (*tsa*), demonstrating the complex nature of the oxidative stress response in this species.

Stress responses other than oxidative were revealed by transcriptome data from *H. parainfluenzae* in coculture, likely due to H_2_O_2_–related damage by *S. mitis*. There was a 2.6-fold increase in expression of the *hfq* chaperone encoding gene, a 2.7-fold increase in Universal stress protein E (*uspE*) and a 2-fold decrease in the repressor *lexA*. LexA is involved in the repression of genes involved in the SOS response of *E. coli* (Kamenšek et al. 2010). Hfq is known to be involved in the stress responses of many species (Deng et al. 2016; Fantappiè et al. 2009). Paralogs of the Universal stress proteins including *uspE*, are known to be involved in response to DNA damage (Gustavsson, Diez, and Nyström 2002). Genes likely involved in DNA repair are also induced in coculture including those encoding DNA ligase (5.4-fold), exodeoxyribonuclease V beta chain (2.4-fold) and endonuclease V (2.7).

*Streptococcus spp*. are known to rapidly consume carbohydrates, and transcriptional data suggest that *H. parainfluenzae* in coculture switches from carbohydrate consumption to alternative sources of carbon and energy. There was increased expression of genes suggesting the breakdown of glycerophospholipids resulting in the uptake and utilization of glycerol, including the predicted extracellular patatin-like phospholipase (2.5-fold), lysophospholipase L2 (3.8-fold), glycerol uptake facilitator protein (5.3-fold), glycerol kinase (3.4-fold), and a fatty acid degradation regulator (2.1-fold). Additionally, there was an increase in expression of fructose 1,6 bisphosphatase (2.8-fold), indicating an active gluconeogenesis pathway, consistent with *H. parainfluenzae* growth on 3-carbon intermediates such as glycerol. There was also evidence of the uptake and catabolism of the sialic acid, N-acetylneuraminic acid as suggested by an increase in expression of SHS family sialic acid transporter (2-fold), sialic acid utilization regulator (3.6-fold), N-acetylneuraminate lyase (2-fold), N-acetylmannosamine kinase (3.8-fold), and N-acetylmannosamine-6-phosphate 2-epimerase (3.1-fold). Lastly, there was an increase in the expression of genes involved in the oligopeptide transport system, *oppA* (3-fold), *oppB* (2.2-fold), *oppC* (2.3-fold), *oppD* (2.7-fold), and *oppF* (2.4-fold). These data together suggest uptake of alternate carbon and energy sources in coculture.

### *In vivo* transcriptional responses of *H. parainfluenzae* vs *in vitro*

We hypothesized that *in vivo* gene expression of *H. parainfluenzae* is substantially impacted by *S. mitis* due to their *in vivo* proximity (Fig. 2) and that our *in vitro* coculture data may be predictive of *H. parainfluenzae* behavior *in vivo*. To quantify this, we compared our *in vitro* monoculture *H. parainfluenzae* transcriptomes to two separate *in vivo* metatranscriptome datasets from dental plaque (Espinoza et al. 2018; Jorth et al. 2014). We aligned each metatranscriptome to the *H. parainfluenzae* genome to generate a dataset of *H. parainfluenzae* transcription within the complex plaque biofilm. Espinoza et al. (2018) quantified meta-transcriptomes of supragingival plaque in health vs disease, and when we compared the healthy samples to our *in vitro* monoculture we observed differential expression of 496 *H. parainfluenzae* genes greater than 2-fold. We then compared these genes to our *in vitro S. mitis* coculture revealing an overlap of 22 genes that were upregulated in both conditions compared to monoculture (Fig. 6A), suggesting that the expression of these genes *in vivo* is due to interactions with *S. mitis*. Interestingly, genes involved in glycerol catabolism and gluconeogenesis were upregulated *in vivo* and in *in vitro* coculture (Table S2). Among the 40 genes downregulated *in vivo* and in *in vitro* cocultures was *ccp* as well as genes involved in methionine metabolism, stringent response, metal transport genes, and *fur* (Table S3). Repeating the same process with the Jorth et al. (2014) metatranscriptome dataset revealed an overlap of 90 upregulated genes (Fig. 6A). Again, this included genes involved in glycerol catabolism and gluconeogenesis shared between all *in vitro* coculture and *in vivo* conditions. These upregulated genes also included several involved in cell stress, DNA damage/repair related pathways, and peptide/oligopeptide transport (Table S4). Among the 55 genes that were downregulated in this comparison were genes known to be involved in oxidative stress including *trxA, tsa*, and again genes involved in methionine metabolism and the stringent response (Table S5). Comparing gene expression patterns shared between all 3 datasets (*in vitro* coculture and both *in vivo* metatranscriptomes) we observed that 18 genes were mutually upregulated including genes involved in glycerol catabolism, gluconeogenesis, and pili biogenesis (Table S6). We also observed 22 genes that were mutually downregulated including those involved in stringent response, methionine metabolism, and *fur* (Table S7, Fig. 6B).

**Figure 6:**
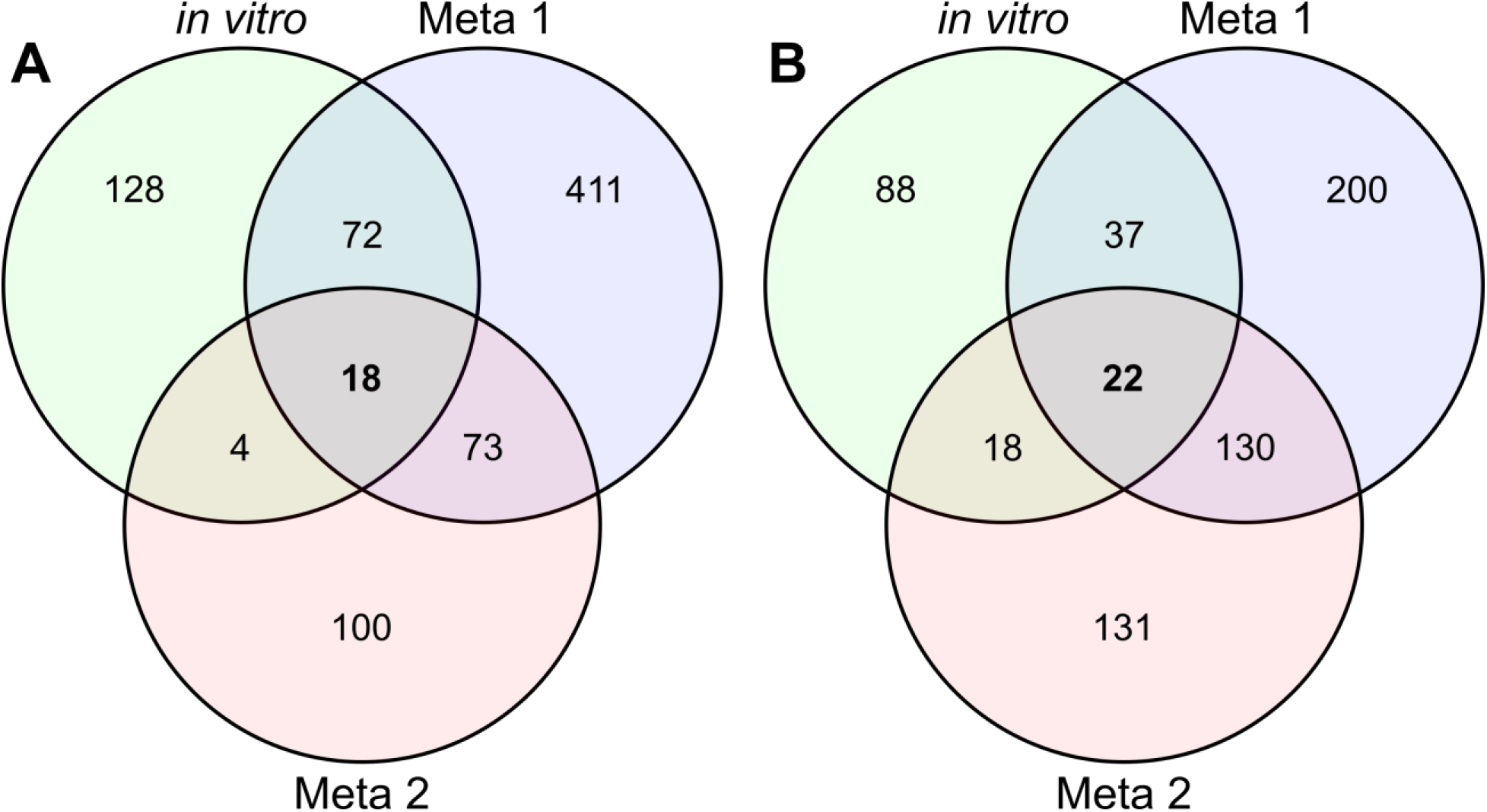
Comparison of coculture gene expression with meta-transcriptome datasets. Our mono vs coculture RNASeq results (*in vitro*) for *H. parainfluenzae* indicated 387 significantly differentially expressed genes (DEG) above 2-fold. By comparing *in vitro* monoculture to *in vivo* metatranscriptome reads from Espinosa 2018 (Meta 1) and Jorth 2014 (Meta 2) we were able to generate two sets of *in vivo*-specific DEGs to determine differences in (A) significantly upregulated or (B) significantly downregulated genes shared between *in vitro* coculture and *in vivo* conditions.

These data suggest that some aspects of *H. parainfluenzae* transcriptional response in the complex biofilms found in plaque can be recapitulated in an *in vitro* coculture with *S. mitis*, which includes a shift away from hexose sugar metabolism. One notably absent overlap between *in vitro* cocultures vs either *in vivo* dataset was the upregulation of genes involved in lactate oxidation observed *in vivo*. Since most *Streptococcus sp*. produce lactate as a metabolic end product (Kreth, Merritt, and Qi 2009), many oral taxa have evolved the ability to utilize lactate as a carbon source. This suggests that *H. parainfluenzae* metabolizes lactate *in vivo*, which it does not do in coculture indicating that further carbon source competition and crossfeeding is likely occurring *in vivo* that did not occur in our *in vitro* coculture on complex medium. In summary, our hypothesis was not borne out in that only a small subset of genes were upregulated in both the co-culture and the *ex vivo* sample. However, the analysis pinpointed interesting commonalities (use of more complex energy sources) and differences (evidence for more complex cross-feeding in the *ex vivo* samples).

## Discussion

Naturally occurring biofilms are often incredibly diverse, complex polymicrobial communities. One such biofilm is human supragingival plaque (SUPP) attached to the tooth surface. This community serves as an excellent site for the study of biofilms and bacterial interactions as its composition and structure are well defined and the majority of species found in this community are cultivable with many being genetically tractable. We chose to study interactions within healthy host microbial communities to gain a better understanding of them which may lead to methods on how to preserve the structure and composition of these communities to prevent dysbiosis and/or colonization by external pathogens. While the behavior of individual species *in vivo* can be inferred from metatranscriptome data we know little about which species might influence one another or by what mechanism(s). Here we detail a reductionist study of one series of interactions between prominent healthy SUPP bacterial species that has potential relevance to multiple host sites.

Both *Haemophilus parainfluenzae* and Mitis group streptococci are highly abundant and prevalent species within the healthy human oral microbiota, especially in SUPP (Eren et al. 2014; Human Microbiome Project Consortium 2012), where we show positive correlations between them (Fig. 1). We chose *H. parainfluenzae* and *Streptococcus mitis* to use in a reductionist approach to ascertain their *in vitro* interactions to compare to *in vivo*. One concern was that while these species coexist in SUPP it is unknown if they exist in close enough proximity for interaction. Our findings demonstrated that these taxa are often found directly adjacent *in vivo* (Fig. 2) and that streptococci seemingly exclude *H. parainfluenzae* above a certain density. This density dependent exclusion also mimicked our *in vitro* findings (Fig. 3) revealing its dependence on *S. mitis* H_2_O_2_ production.

Streptococci H_2_O_2_ production is thought to provide a competitive advantage *in vivo* and presents a source of stress that coexisting bacterial species must tolerate. We and others (Redanz et al. 2018; L. Zhu and Kreth 2012; B. Zhu et al. 2019; L. Zhu and Kreth 2012; Liu et al. 2011) have previously shown potential adaptive responses of bacterial species to streptococci-produced H_2_O_2_ and investigated *H. parainfluenzae’s* as well (Fig. S1). We discovered a multifactorial, highly redundant oxidative stress response that differs from other closely related species, particularly *H. influenzae*. *H. parainfluenzae* possesses *oxyR* whose gene product upregulates expression of genes that encode catalase and other H_2_O_2_ resistance related proteins. We demonstrated that while loss of OxyR caused a significant increase in H_2_O_2_ sensitivity, loss of catalase or other individual gene products that often provide H_2_O_2_ resistance did not (Fig. 4A) which directly contrasts the significant importance of catalase in *H. influenzae*, whose deletion leads to its inability to survive high concentrations of H_2_O_2_ (Juneau et al. 2015; Bishai et al. 1994). We also observed that impaired H_2_O_2_ stress responses by *H. parainfluenzae* led to a decrease in fitness of *S. mitis* in *in vitro* coculture (Fig. 4B) and when *H. parainfluenzae* H_2_O_2_ detoxification was substituted with exogenous catalase we also saw a considerable growth benefit to *S. mitis* (Fig. S2). However, we also observed that *H. parainfluenzae* still further increased *S. mitis* yield, even when exogenous catalase was present. These and our *in vivo* microscopy data suggest further mutual benefit between these species beyond H_2_O_2_ responses.

Unexpectedly, *H. parainfluenzae* was unable to obtain enough NAD to grow from host saliva which bathes the SUPP environment where it resides. We demonstrated that abundant oral *Corynebacterium spp*. were also unable to complement *H. parainfluenzae’s* NAD auxotrophy yet Mitis group streptococci could do so (Fig. 5). Given their close proximity in SUPP and overlap in other body sites, *Streptococcus sp.-*produced NAD could therefore be an important determinant of *H. parainfluenzae’s* ability to survive and colonize various sites of the human body. It is interesting to note that both *H. parainfluenzae* and *S. mitis* are found as commensals not just in the same sites of the human oral cavity (Mark Welch, Dewhirst, and Borisy 2019), but also in other sites in the nasopharynx (Könönen et al. 2002; Kosikowska et al. 2016). When cocultured with *S. mitis in vitro, H. parainfluenzae* transcriptomes indicated a shift from carbohydrate consumption to alternative sources of carbon and energy including glycerophospholipids, sialic acid, oligopeptides, and the initiation of gluconeogenesis, consistent with growth on short carbon chain compounds. Downregulation in coculture of genes involved in the stringent response also suggest new access to peptides. When comparing the *H. parainfluenzae* coculture *in vitro* transcriptome to *in vivo* SUPP metatranscriptomes (Fig. 8) we observed similar regulation of these same carbon source pathways. These data coupled to our *in vivo* observations of direct proximity, strongly suggest that *S. mitis* induces these same behaviors in *H. parainfluenzae in vivo* that we observed *in vivo* and highlights the utility of this reductionist mechanistic study. One notable pathway absent from our *in vitro* cocultures was lactate oxidation which has previously been shown to be critical for co-infection between organisms with *Streptococcus spp*. by cross-feeding on this fermentation product (Ramsey, Rumbaugh, and Whiteley 2011). *In vivo*, metatranscriptome data indicated that *H. parainfluenzae* was also upregulating lactate oxidation gene products which would be expected in the more diverse, competitive SUPP environment compared to *in vitro* coculture alone.

Currently, there is a wealth of information and techniques available to study the composition and structure and overall gene expression of complex naturally occurring biofilms. However, specific mechanisms of interaction within these complex environments remain elusive and are worthy of study as a means to identify potential routes to keep these complex communities intact which may benefit associated host(s) or the environment. We chose two highly abundant and prevalent species within the human supragingival plaque biofilm and discovered mechanisms responsible for their micron-scale distribution within this environment while also revealing additional factors at play in the *in vivo* community. This study demonstrates that *H. parainfluenzae* co-localizes with Mitis group streptococci in supragingival plaque and that this co-existence is dependent on the relative densities of each taxon. We demonstrate that this density-dependent exclusion of *H. parainfluenzae* is due to the production of H_2_O_2_. The response of *H. parainfluenzae* to H_2_O_2_ appears unique compared to other *Haemophilus sp*. responses as a more redundant system where catalase and other gene products provide an equally modest level of protection. *H. parainfluenzae* H_2_O_2_ detoxification supports the growth of *S. mitis*, while *S. mitis* supports the growth of *H. parainfluenzae* by providing NAD. While our work focuses on supragingival plaque, these observations likely translate to other aerobic sites where these organisms frequently overlap. We report here for the first time several mechanisms that underlie the coexistence between these two species which are highly abundant and prevalent in the human host as part of a diverse biofilm to provide a starting point for further study of this community and its relevance to host health.

## Materials and Methods

### Strains and Media

Strains and plasmids used in this study are listed in Table 1. Unless indicated, *Streptococcus mitis* was cultured using Brain Heart Infusion (BHI) broth or solid agar supplemented with Yeast extract (YE), *Haemophilus parainfluenzae* had additional supplementation with 14μg/ml Nicotinamide Adenine Dinucleotide (Sigma-Aldrich) and 14μg/ml Hemin (Sigma-Aldrich) - (BHI-YE-HP). *Escherichia coli* was grown on Luria Broth (LB). *H. parainfluenzae* and *S. mitis* were grown at 37°C and 5% CO_2_, *E. coli* at 37°C in standard atmospheric conditions with liquid cultures shaken at 200 RPM. Antibiotics were used at the following concentrations: Kanamycin 40μg/ml, Vancomycin 5μg/ml, and Spectinomycin 50μg/ml for *E. coli*, 200μg/ml for *H. parainfluenzae*.

**Table 1:**
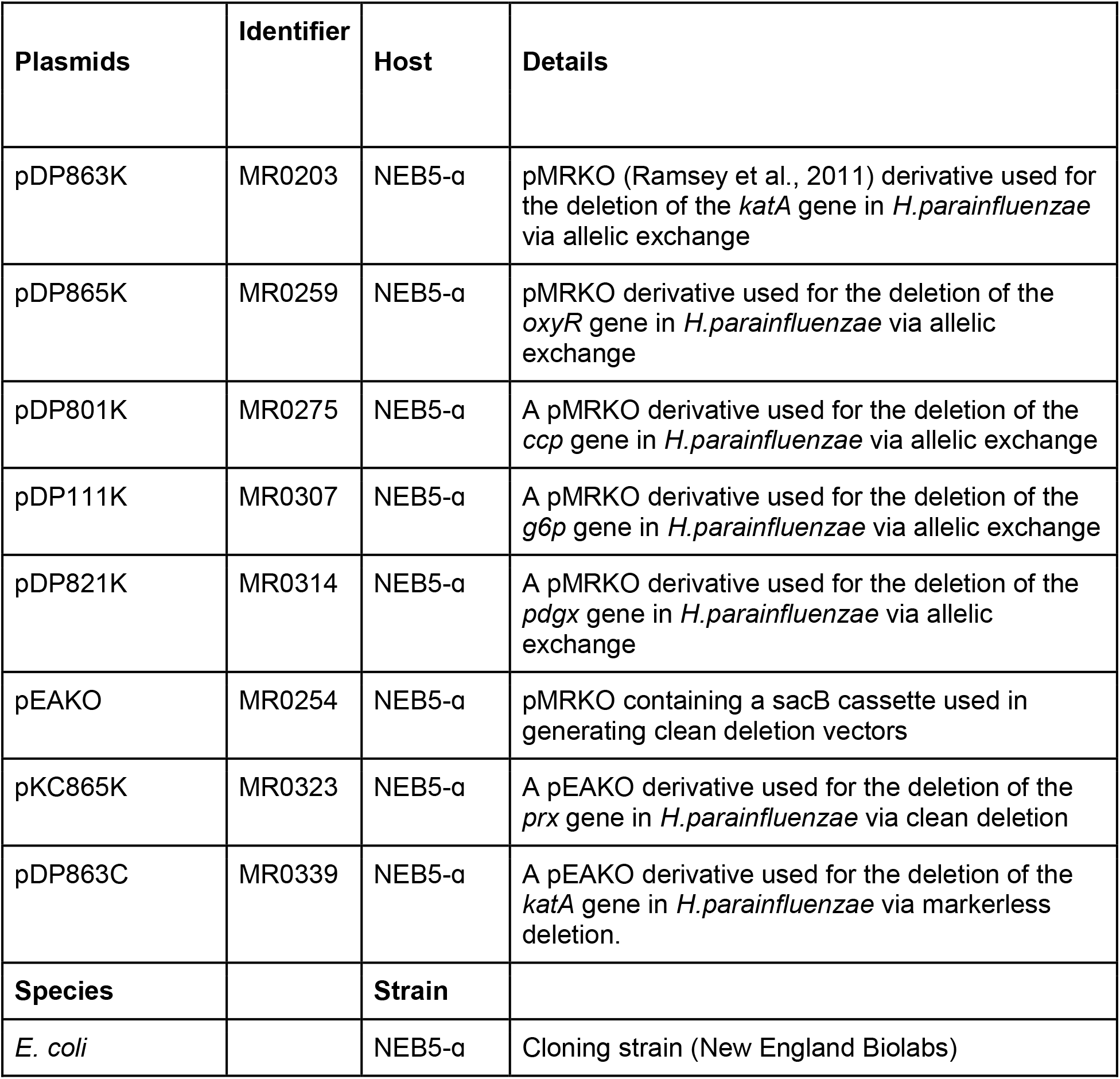
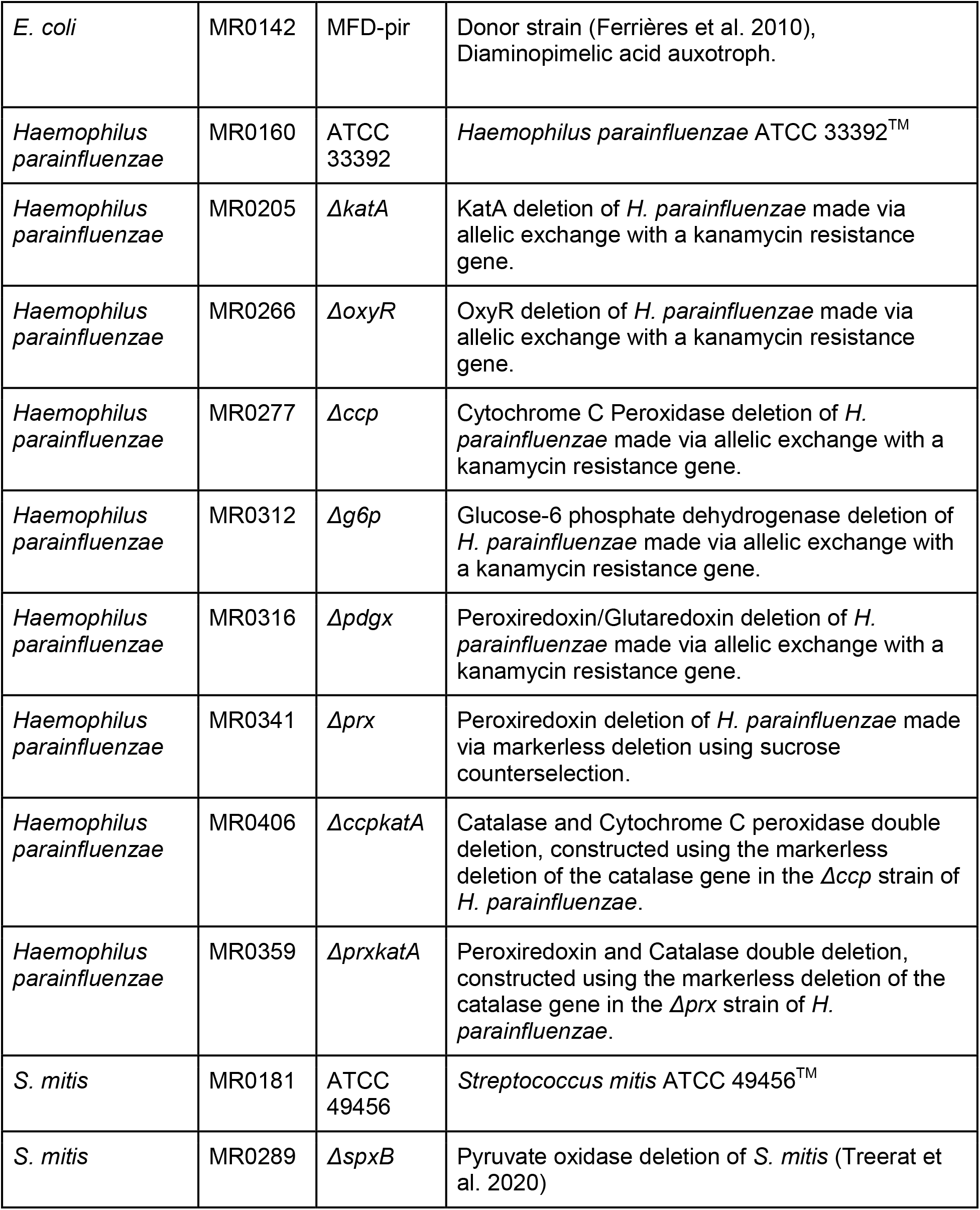
Strains and plasmids used in this study

**Table 2:**
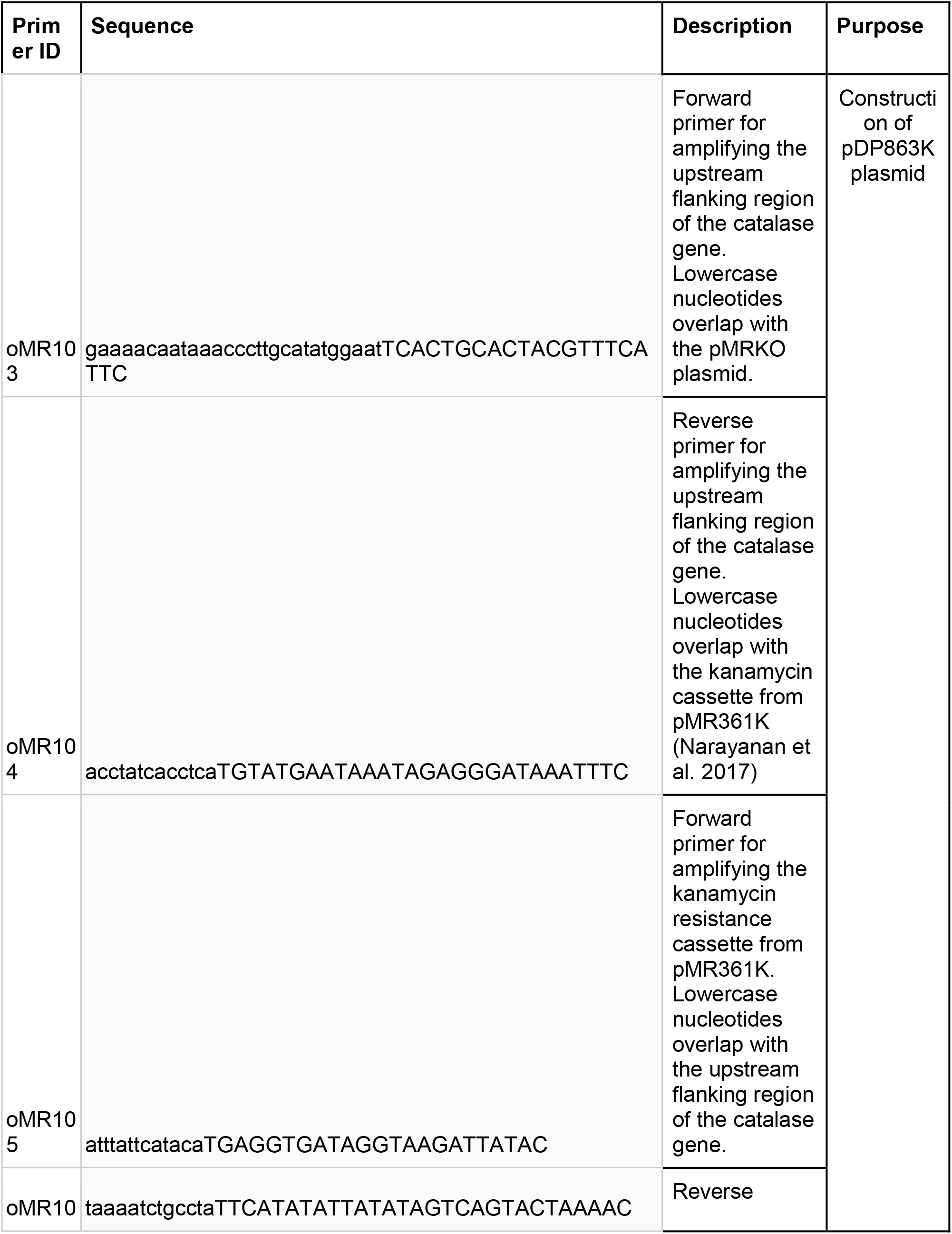
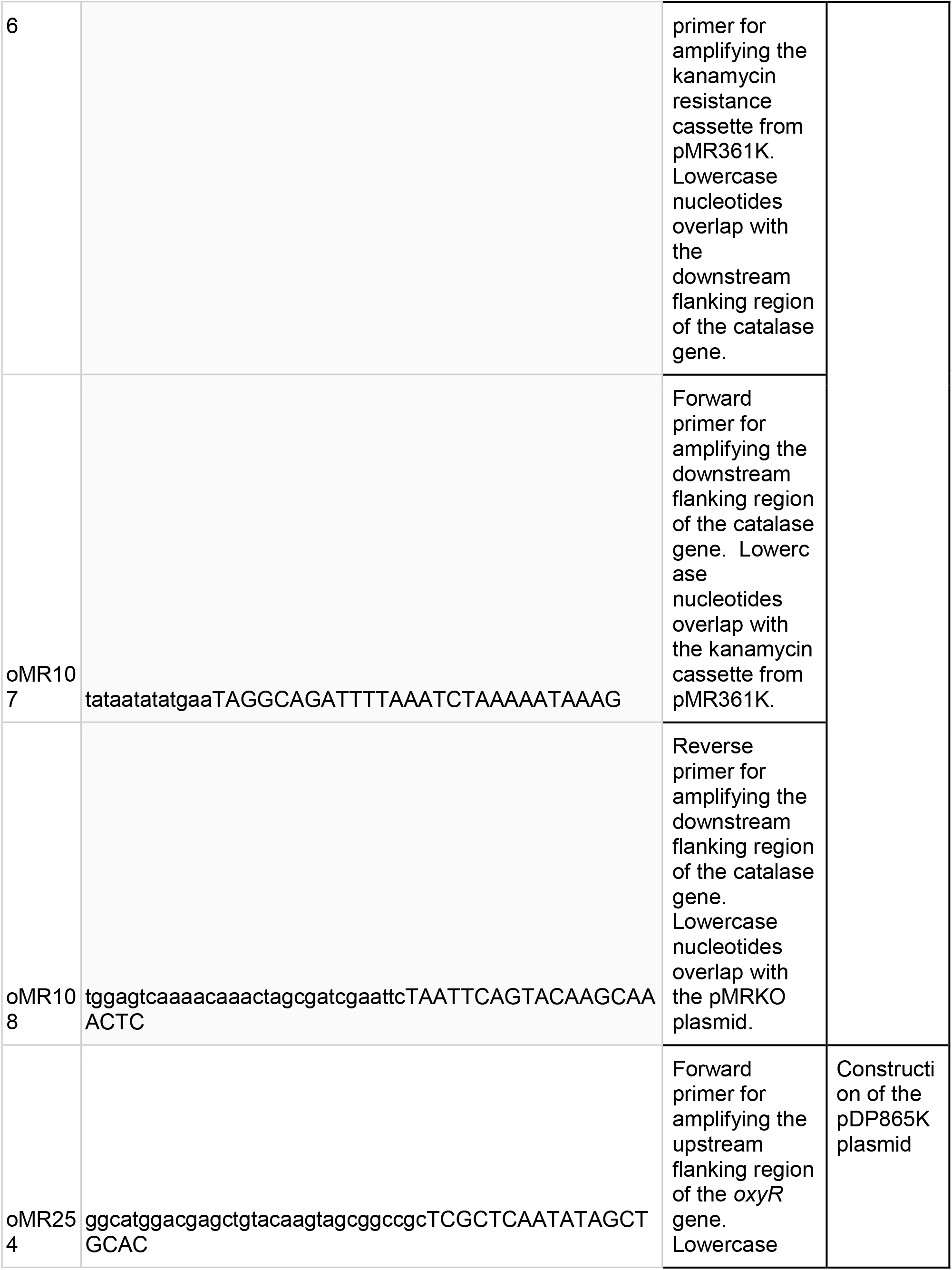
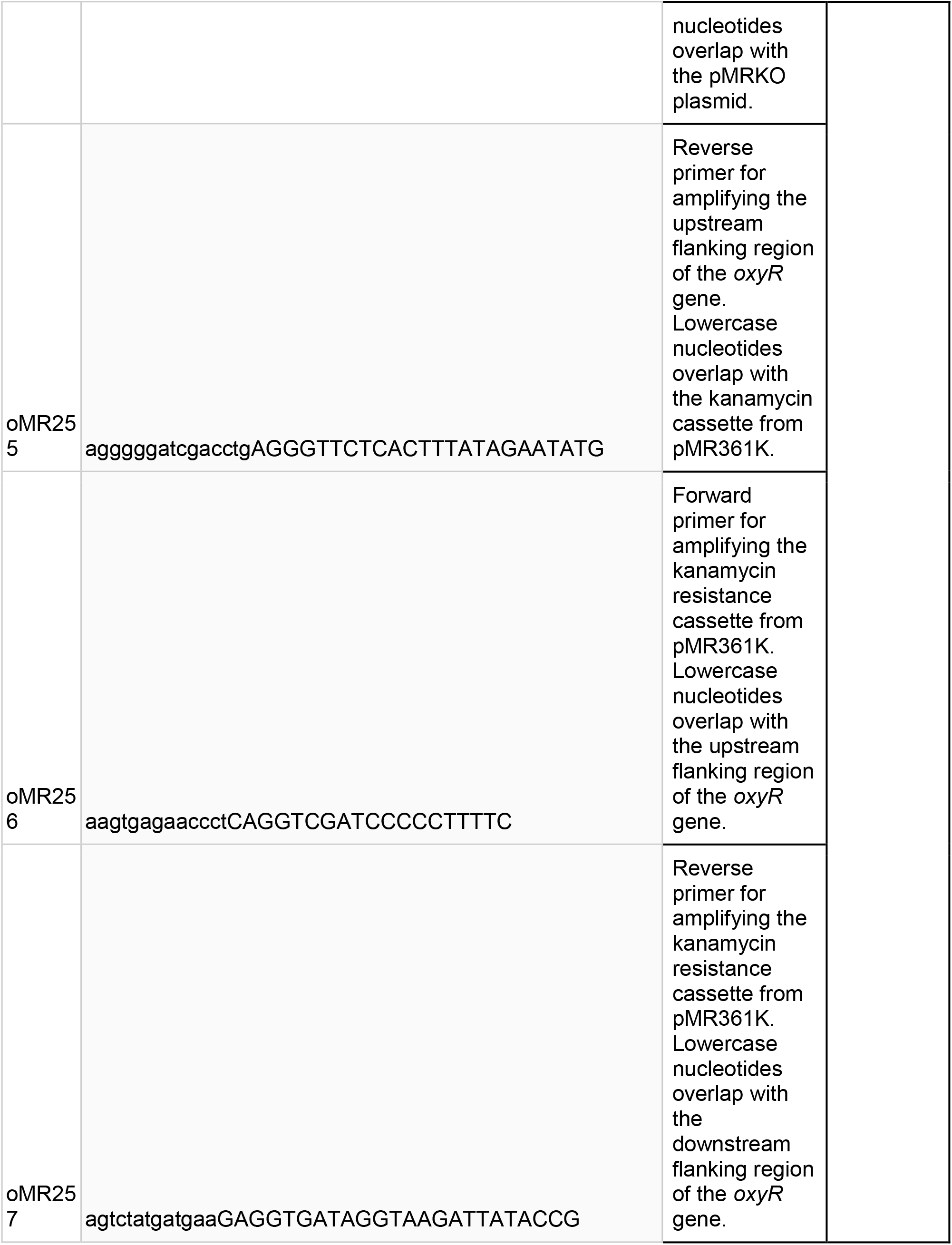
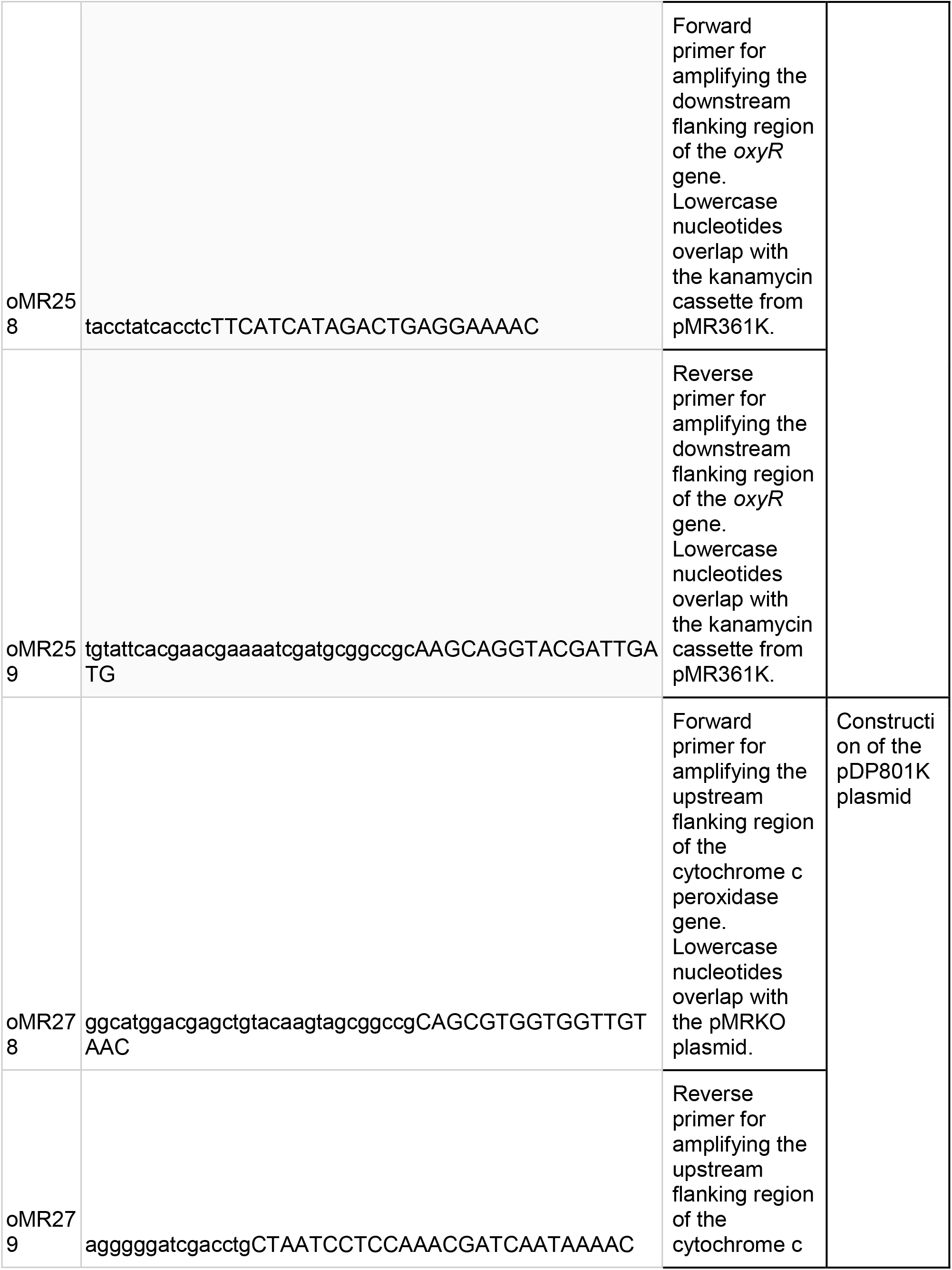
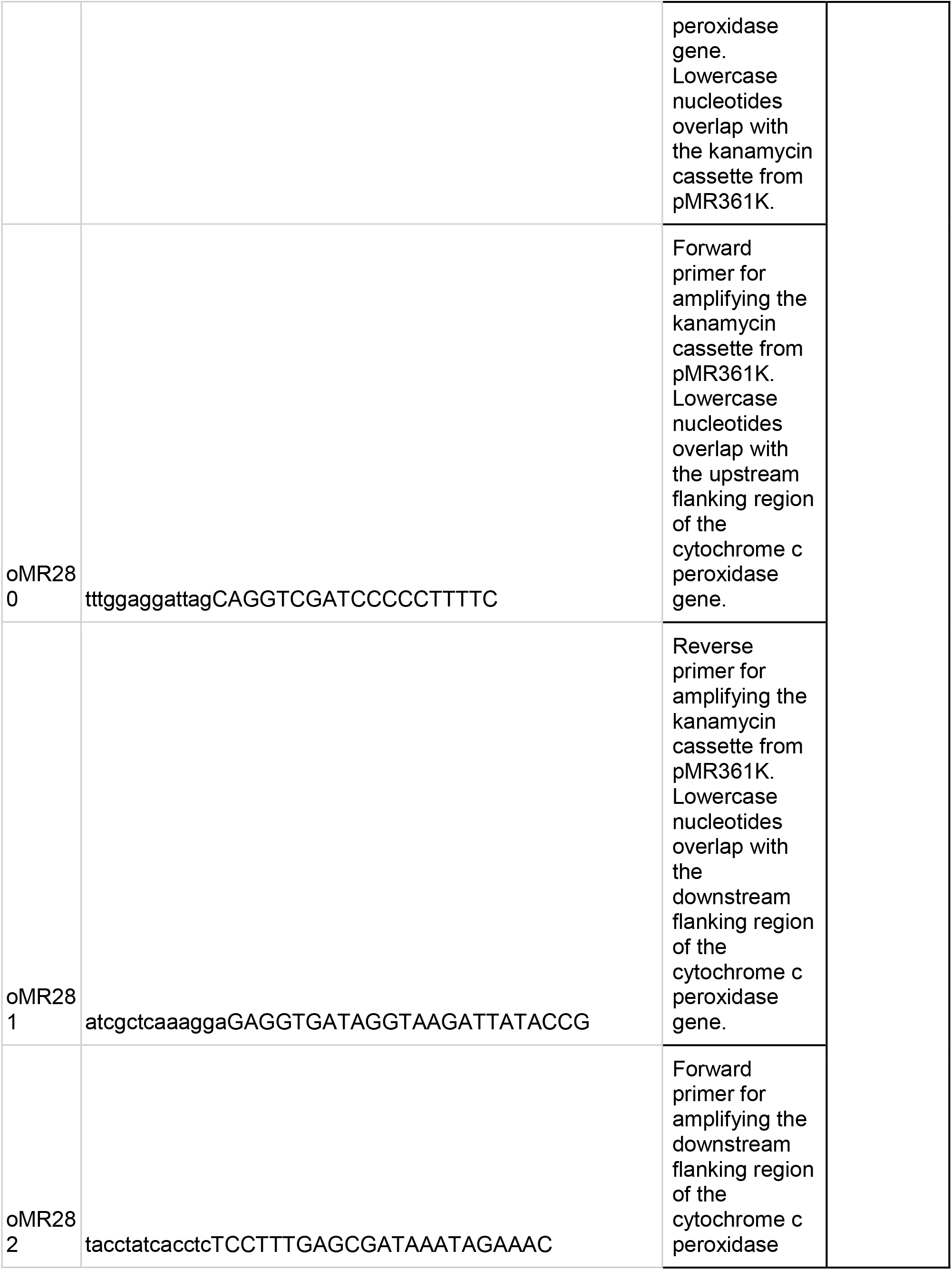
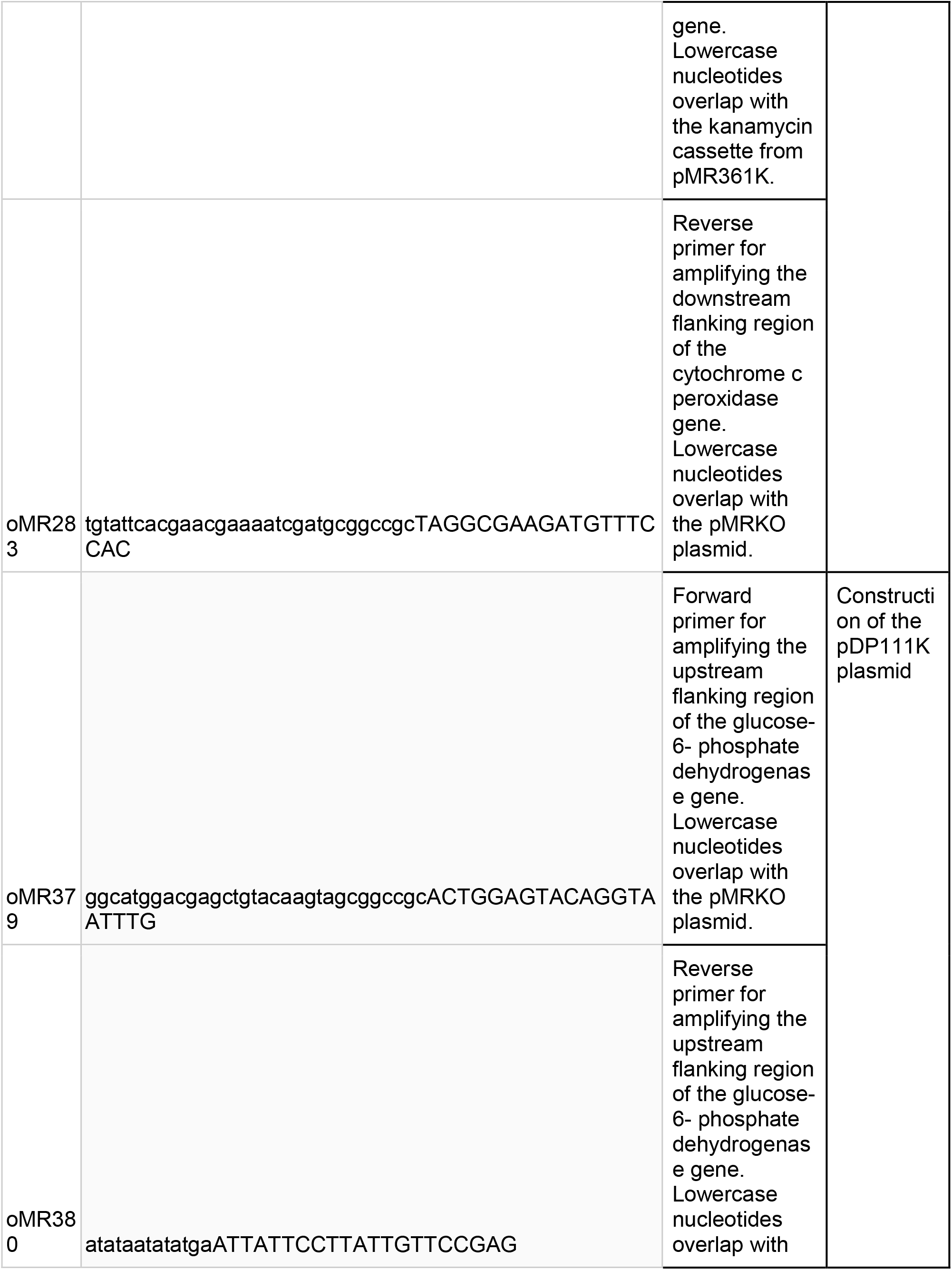
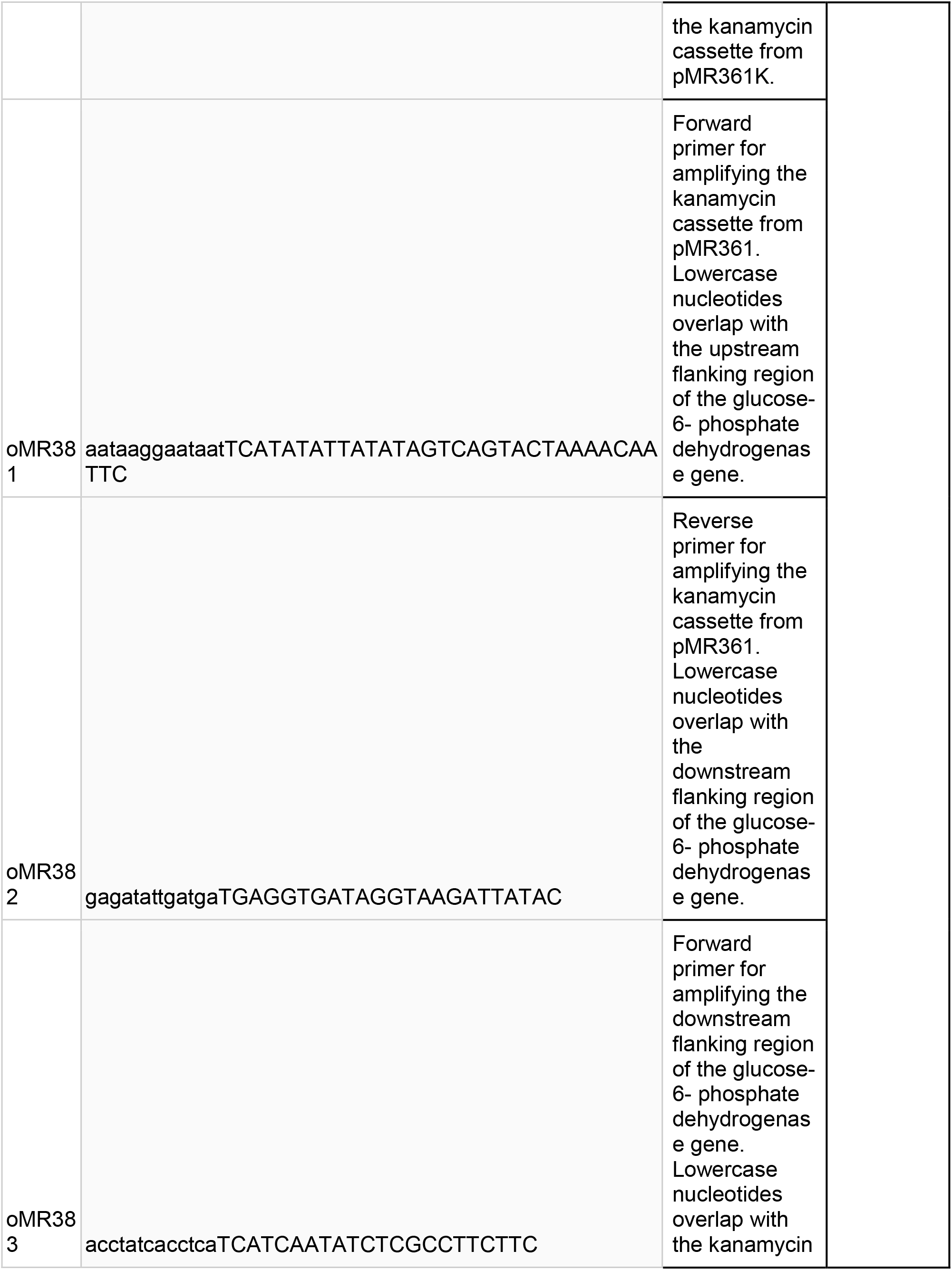
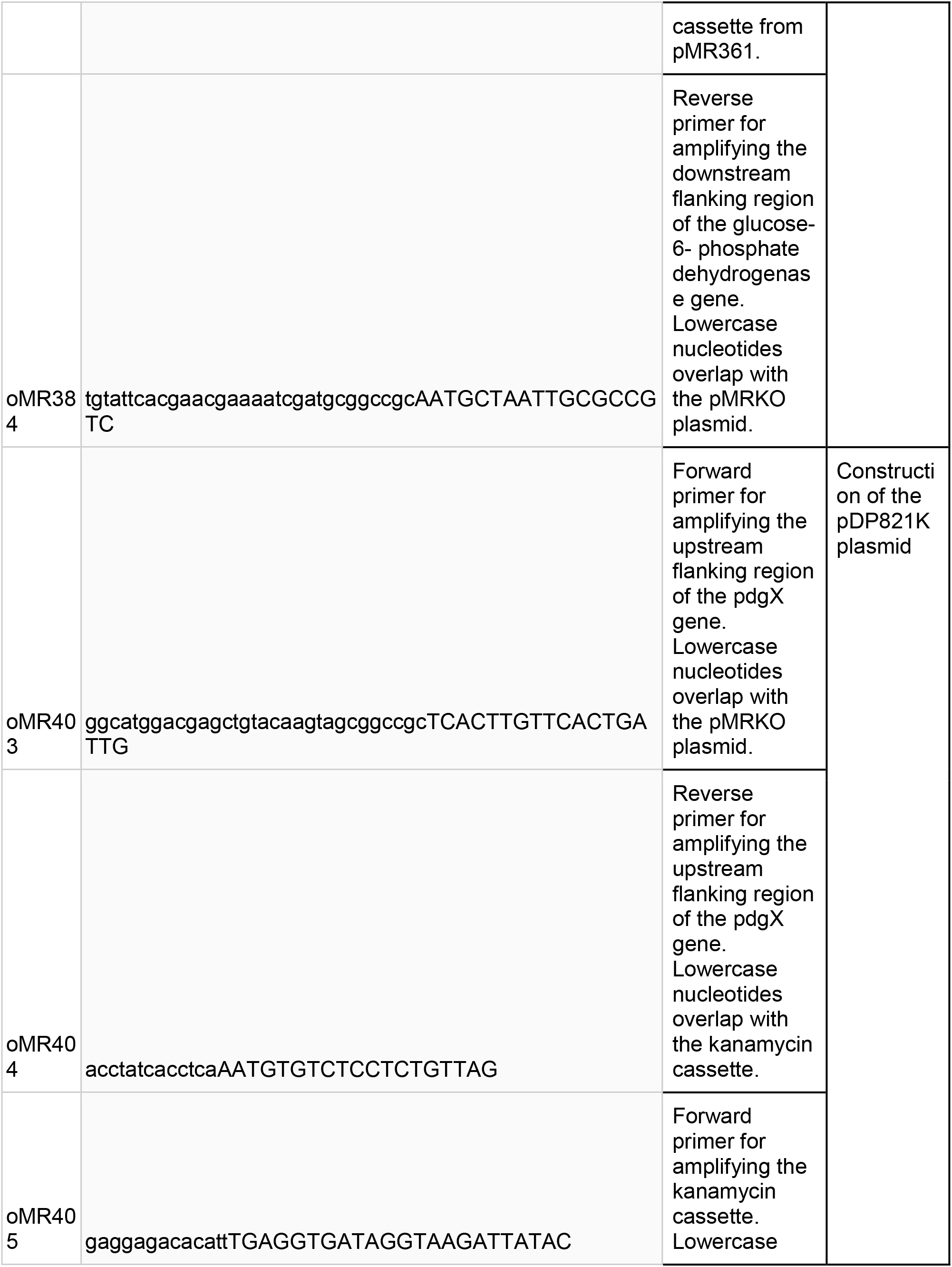
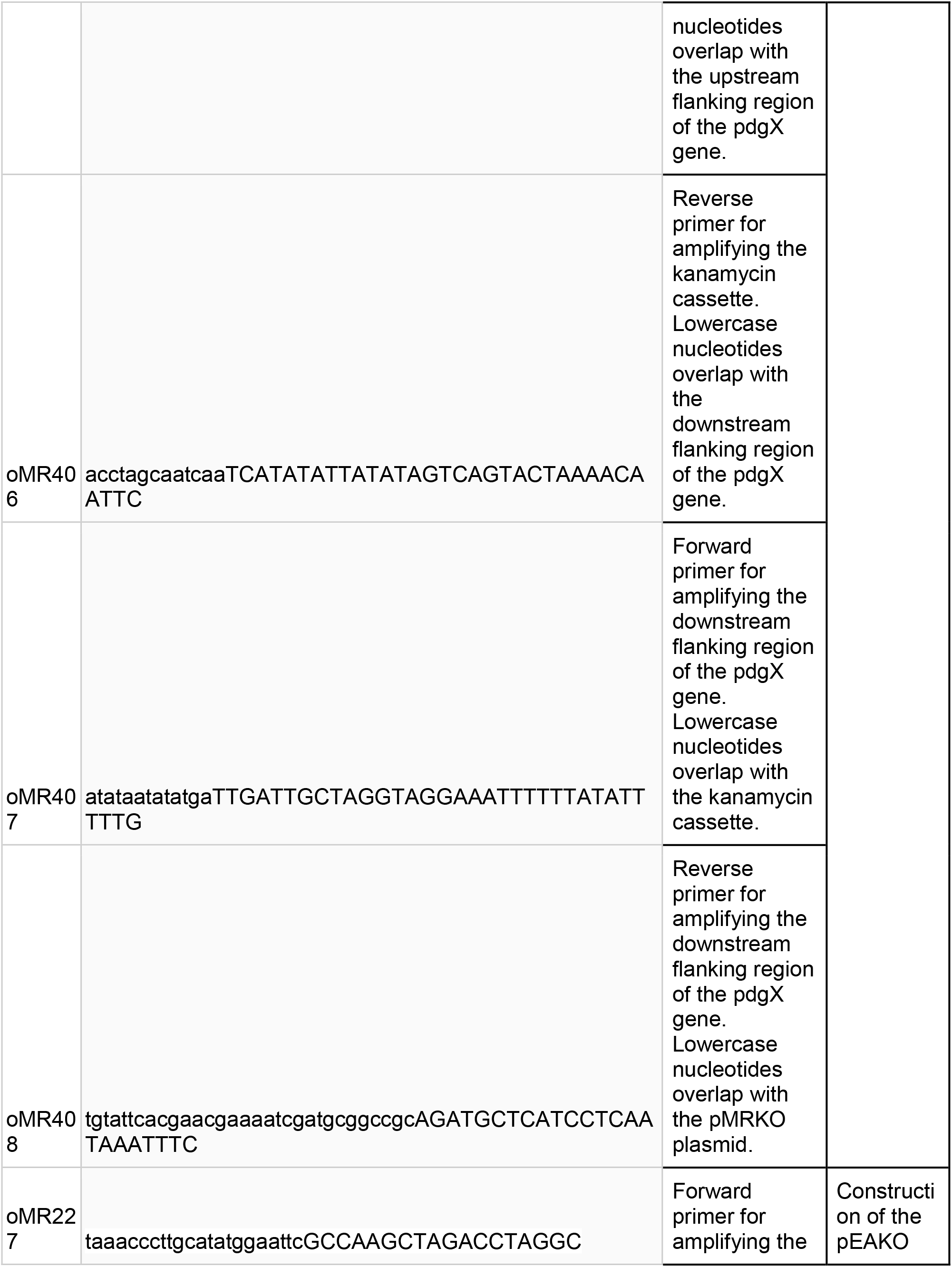
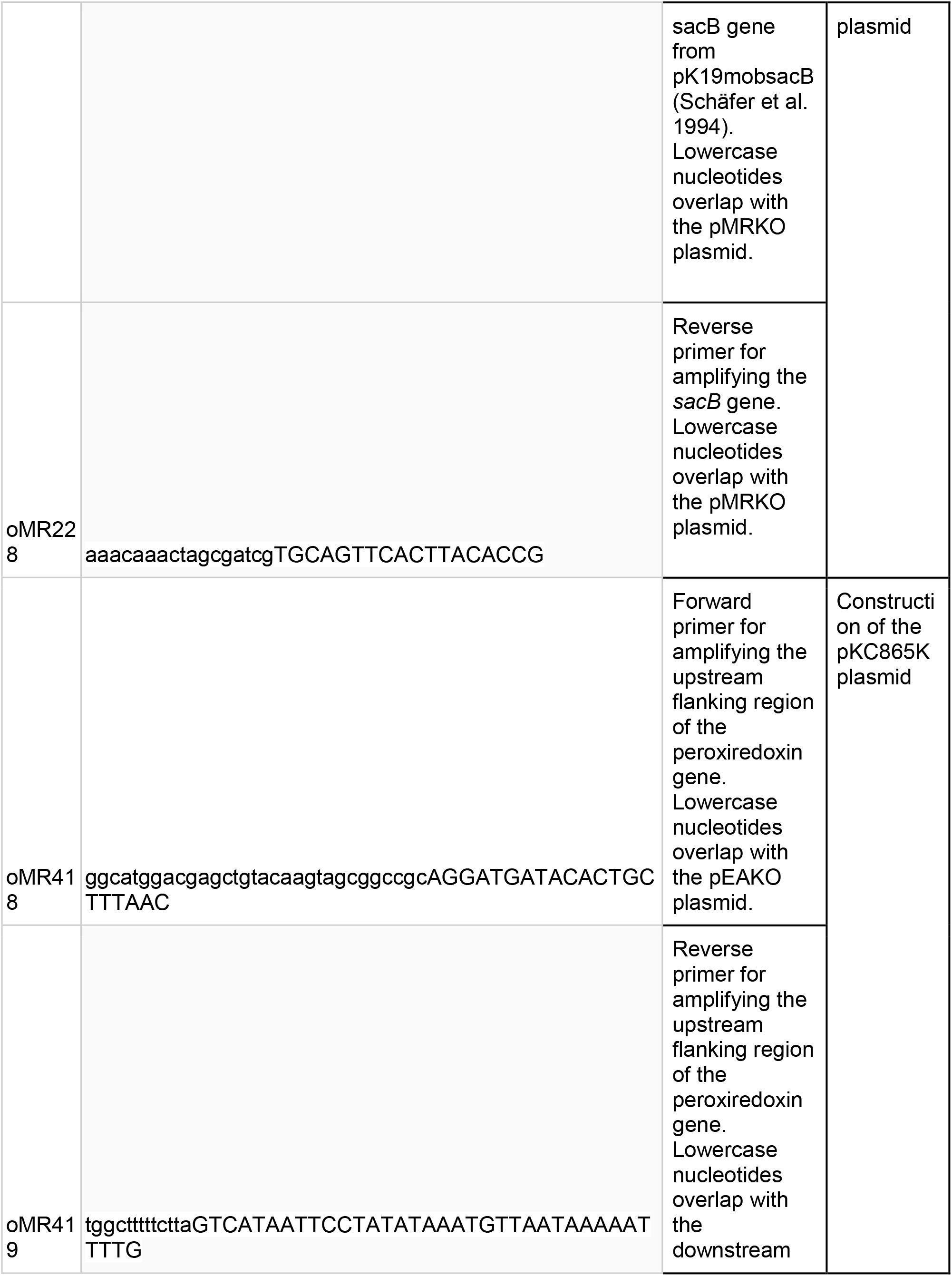
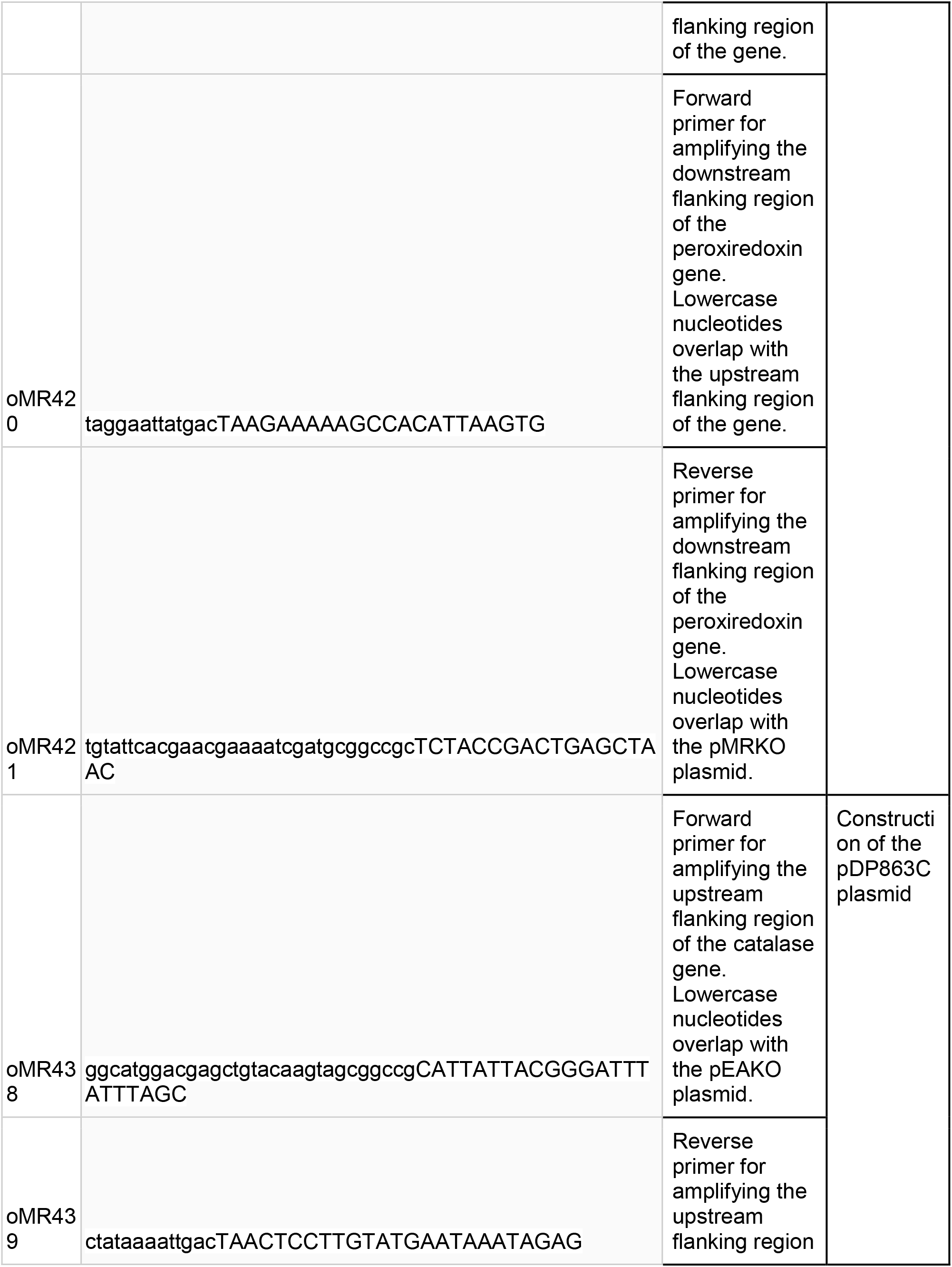
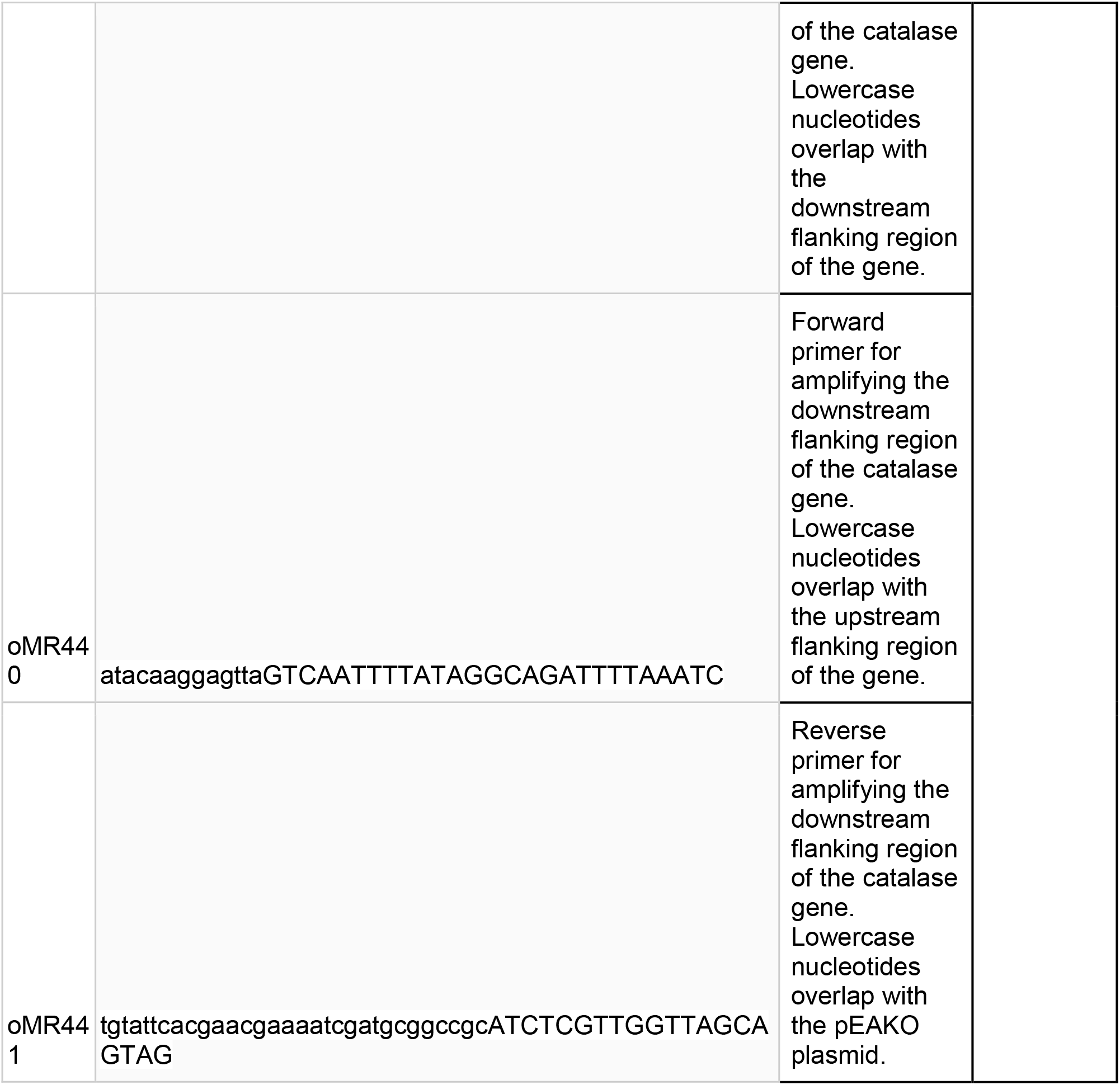
Primers used in this study

**Table 3:**
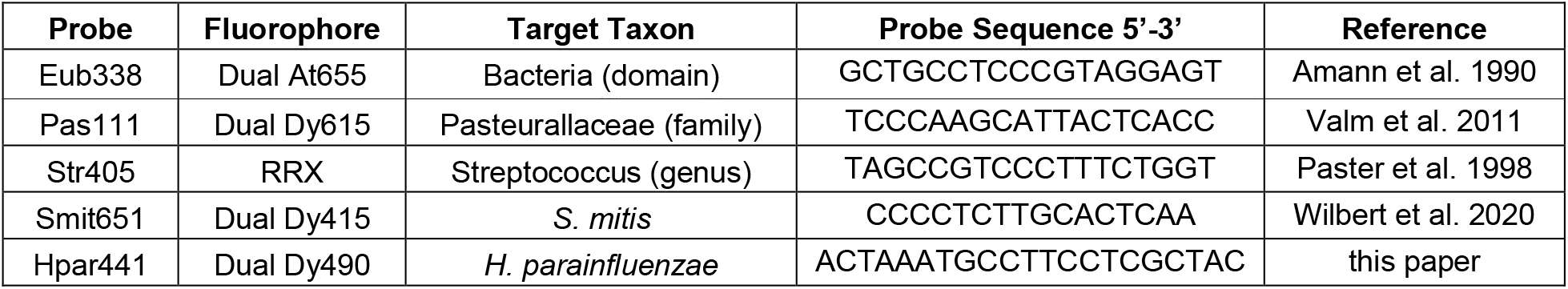
DNA FISH probes

### Genomic and plasmid DNA isolation

*H. parainfluenzae* Genomic DNA was isolated using the DNeasy Blood & Tissue kit (Qiagen) according to the manufacturer’s instructions. Plasmid isolations were performed using QIAprep spin miniprep kits (Qiagen).

### Genetic manipulation of *H. parainfluenzae*

Gene deletions were generated using derivatives of a suicide vector pMRKO (Ramsey, Rumbaugh, and Whiteley 2011). 1kb flanking regions of the target gene were amplified via PCR using Q5 High-fidelity 2X Mastermix and primers indicated in Table S5. For allelic exchange, these fragments were assembled to flank a kanamycin resistance cassette via isothermal assembly using the NEBuilder HiFi DNA Assembly MasterMix (New England Biolabs) and cloned into pMRKO. This resulting reaction was then transformed into NEB5α competent cells using the manufacturer’s instructions (New England Biolabs). Plasmid constructs were verified via restriction digests and Sanger sequencing. After screening, plasmids were transformed into the donor strain MFD-pir, using the TSS transformation method (Chung, Niemela, and Miller 1989). These strains were then used to conjugate into *H. parainfluenzae*. Briefly, washed cells of *H. parainfluenzae* overnight cultures were subjected to heat-shock (46°C for 6 minutes) and combined with the donor strain by spread plating on a BHI-YE HP agar plate supplemented with 0.3 mM di-amino pimelate (DAP). Plates were then incubated overnight at 37°C in 5% CO_2_. Cells were then harvested and dilutions were plated on BHI-YE HP with 40 μg/ml of kanamycin and incubated for 24-48 hours at 37°C in 5% CO_2_. Mutants were then screened by testing for sensitivity to spectinomycin (spectinomycin resistance cassette on pMRKO backbone), PCR and Sanger sequencing.

The markerless deletion of genes in *H.parainfluenzae* involved modifications to the above protocol. 1kb flanking regions were amplified and cloned into a pMRKO derivative containing a *sacB* gene (pEAKO - Table 1). Plasmids were then transformed into *H. parainfluenzae* via conjugation as described above. After transformation, cells were subjected to counterselection by plating on BHI-YE HP containing 10% sucrose for 4-5 days. Mutants were then screened via PCR and Sanger sequencing.

### Read abundance data

MetaPhlAn (Segata et al. 2012) species-assigned metagenomic sequence data from the Human Microbiome Project (Human Microbiome Project Consortium 2012) for the “Supragingival Plaque” oral site was 1^st^ sorted based on predicted read abundance for *Haemophilus parainfluenzae*. Using the top quartile (highest 25% of samples enriched for *H. parainfluenzae*) we compared these samples to the bottom 75% and performed LEfSe analysis (Segata et al. 2011) to predict species likely to be significantly encountered at higher *H. parainfluenzae* abundance. LDA scores of a log10 score of ≥3 were deemed significant.

### Plaque Collection, Fixation, and Storage

We collected supragingival plaque samples by using a toothpick to scrape the surface of 7 donors’ teeth, avoiding the gingival margin. These donors were instructed to refrain from practicing oral hygiene for 24 h before collection. We fixed the plaque in a solution of 2% paraformaldehyde (PFA) in PBS buffer on ice for 2 to 6 h. The PFA was removed by three washes with 10 mM Tris HCl buffer (pH 7.5). The samples were stored in a 1:1 (vol/vol) solution of 10 mM Tris HCl (pH 7.5) and 100% ethanol at –20°C.

### DNA FISH and Mounting

FISH with 16S DNA probes was conducted on the surface of UltraStick Slides (Thermo Scientific). Aliquots of plaque were dried on the slides for 10 min at 46°C. The plaque was hybridized with 2 μM of each probe in a 900 mM NaCl, 20 mM Tris HCl (pH 7.5), 0.01% SDS, and 20% formamide hybridization buffer for 3 h in a humid chamber at 46°C. Non-hybridized probe was removed by washing the slides in prewarmed 215 mM NaCl, 20 mM Tris HCl (pH 7.5), and 5 mM EDTA wash buffer for 15 min at 48°C. The slides were rocked once during the wash incubation. The slides were rinsed in chilled deionized water and allowed to mostly air-dry before the samples were mounted in ProLong Gold Antifade mounting medium (ThermoFisher) under a #1.5 coverslip. The slides were dried flat in the dark.

### Imaging

We imaged the hybridized plaque with an LSM 780 Confocal Microscope (Zeiss) with a Plan-Apochromat 40x/1.4 Oil DIC M27 objective. Each field of view was simultaneously excited by linear scanning with 405, 488, 561, and 633 nm laser lines. The emission spectra for the probes’ fluorophores were decomposed by linear unmixing using ZEN software (Zeiss) using reference spectra recorded from pure *Leptotrichia buccalis* culture samples hybridized with the appropriate fluorophore as described above. To obtain a random sample of the masses of plaque large enough to permit spatial analysis, we scanned transects spaced every 5 μm along the coverslip at 40x magnification and imaged every mass of plaque that was at least 70 μm in diameter and 250 μm away from the previously imaged fields of view. For each donor, we either imaged the first 20 fields of view that satisfied these criteria or as many fields of view as were present on the slide. To maximize the number of bacteria captured in each image, we imaged the focal plane closest to the surface of the slide.

### Image Analysis

To allow quantitative analysis of the spatial distribution of the taxa of interest, we used FIJI to create binarized *S. mitis, H. parainfluenzae*, and bacterial mass images (Schindelin et al. 2012). A slight misalignment of the Smit651 channel was brought into closer alignment with the other channels by shifting it up by 2 pixels, cropping 3 pixels off each edge, and re-scaling the channel image to regain the original 2,048 by 2,048 resolution. The noise in each channel was reduced by applying a median filter with a radius of 3 pixels. To create a bacterial biomass mask, the Eub338 channel was automatically segmented by thresholding with the global Otsu method (Otsu 1979) and dilating the segmented area by 3 pixels. The *S. mitis* channel was created by segmenting the Str405 and Smit651 channels with the local Bernsen and global RenyiEntropy automatic thresholding methods, respectively (BERNSEN 1986; Kapur, Sahoo, and Wong 1985). Both segmented images were combined using the Boolean “AND” operator to retain the pixels appearing in both images. The *H. parainfluenzae* channel was created by segmenting the Pas111 and Hpar441 channels with the local Bernsen and global RenyiEntropy automatic thresholding methods, respectively. Both segmented images were combined using the Boolean “AND” operator. To ensure that there was a sufficiently large area of *H. parainfluenzae* in the images for reliable analysis, only the 41 fields of view in which at least 1% of the bacteria mass was covered by *H. parainfluenzae* in the associated binary image were used for the following analyses.

We evaluated the pair correlations between *S. mitis* and *H. parainfluenzae* over different distances using a linear dipole analysis performed in Daime 2.2 (Daims, Lücker, and Wagner 2006; Daims and Wagner 2011). For this analysis, the reference space in each image was restricted to the area in the binary bacterial biomass image. We used all possible dipoles with lengths ranging from 0.15 to 99.90 μm in steps of 0.45 μm.

We evaluated trends between the local densities of both taxa, by dividing each field of view into 1024 6.64 μm by 6.64 μm blocks, discarding blocks that did not contain any of the binary bacterial mass image, and calculating the fraction of each block that was covered by the *S. mitis* and *H. parainfluenzae* binary images.

### Mono and Coculture Assays

Colony biofilm assays were carried out as described previously (Ramsey et al. 2016). Briefly, equal volumes of *H. parainfluenzae* and/or *S. mitis* were spotted either in mono or coculture on sterile 25mm 0.2 μm polycarbonate membranes (MilliporeSigma™) that were placed on BHIYE HP agar plates following adjustment of optical density. *H. parainfluenzae* was spotted at an OD_600_ of 1 and *S. mitis* at OD_600_ of either 0.1, 1 or 3, with 10μl for each strain. The plates were then incubated for 24 hours at 37°C in 5% CO_2_. The membranes were then transferred to 1ml sterile media, vortexed and pipetted to ensure complete resuspension of the colony into the media; serially diluted and plated for CFU enumeration. *S. mitis* was enumerated by counting CFUs on BHI-YE and *H. parainfluenzae* on BHI-YEHP with 5 μg/ml vancomycin.

### Disk diffusion assays

Cultures of *H. parainfluenzae* were grown anaerobically in BHI-YEHP overnight. All strains were then adjusted to an OD_600_ of 1 and 100μl was spread plated on BHIYE HP plates and incubated aerobically for 2 hours at 37°C in 5% CO_2_. 5 μl of 30% H_2_O_2_ was then added to a sterile 5mm paper disk and plates were incubated for 24 hours at 37°C in 5% CO_2_. The diameters of the zones of inhibition were then measured using a caliper in at least 3 axes.

### Coculture transcriptome sample preparation

RNASeq analyses were carried out on mono and coculture samples following the colony biofilm assays described above. Briefly, *H. parainfluenzae* was spotted on the polycarbonate membranes at an OD_600_ of 1 and *S. mitis* at OD_600_ 0.1, with 10μl for each strain in either mono or coculture. The plates were then incubated for 22 hours at 37°C in 5% CO_2_. The membranes were then transferred onto fresh media for 4 hours and immediately placed in RNAlater solution (Ambion™). Experiments were carried out in biological duplicates. RNA extraction, library preparation and sequencing were then carried out by the Microbial ‘omics core facility at the Broad Institute. Sequences are submitted to the NIH SRA Gene Expression Omnibus (GEO) database. Bioproject number is pending on SRA approval and will be made available upon publication or by request to the corresponding author.

### Transcriptome analyses

Genome data for *H. parainfluenzae* ATCC 33392 and *S. mitis* NCTC 12261 was obtained from NCBI and genome annotations were generated using RAST under default settings (Brettin et al. 2015; Overbeek et al. 2014; Aziz et al. 2008).

RNASeq reads were aligned, mapped and differentially expressed genes were analyzed using bowtie2, HTSeq, DESeq2 and R using a custom Unix and R pipeline (available at https://github.com/dasithperera-hub/RNASeq-analysis-toolkit).

Sequencing reads were aligned to the respective genomes using bowtie version 2.2.4 (Langmead and Salzberg 2012) with the default parameters. Reads aligning to coding sequences were then counted using the HTSeq-count function from HTSeq version 0.8.0 (Anders, Pyl, and Huber 2015) using the “intersection-nonempty” mode. DESeq2 (Love, Huber, and Anders 2014) was then used to carry out normalization and differential expression analysis. Transcripts were considered statistically significant if they had an adjusted p-value of less than 0.05 (FDR<0.05). Genes considered to be overexpressed had a fold change greater than 2, whilst genes under-expressed had a fold change less than 0.5 (−2-fold). Following identification of differentially expressed genes (DEGs), KEGG annotations were obtained for each of the genomes using BlastKOALA (Kanehisa, Sato, and Morishima 2016). Pathway analysis for the DEGs was then carried out by mapping to KEGG orthology (https://www.genome.jp/kegg-bin/get_htext?ko00001). This allowed for improved annotations and the identification of potential pathways that are involved in coculture. The same pipeline was used to analyze *H. parainfluenzae* gene expression in published metatranscriptome datasets (Jorth et al. 2014; Espinoza et al. 2018). For these analyses, gene expression from *H. parainfluenzae* monoculture was compared to *H. parainfluenzae* gene expression in coculture and metatranscriptome datasets.

### Complementing Nicotinamide Adenine Dinucleotide (NAD) auxotrophy of *H. parainfluenzae*

Overnight cultures of *H. parainfluenzae* were washed 3 times in 1x Phosphate buffered saline (PBS) and diluted to an OD_600_ of 0.1 and spread plated on a plate containing BHI-YE supplemented with Hemin and 20units/ml catalase. 5μL of bacteria at an OD_600_ of 1 was added to a sterile paper disk and incubated for 48 hours. Strains used for spotting include *Corynebacterium matruchotii, C. durum, Streptococcus mitis, S. sanguinis, S. cristatus*, and *S. gordonii*.

## Author Contributions

MR and JMW designed research; EA, KC, KCL, AM, VML, and DP performed research, JMW, AM, DP and MR wrote the paper.

## Acknowledgements

We thank Janet Atoyan and the RI-EPSCOR sequencing facility at URI for sequence generation, Jonathan Livny and the Microbial ‘Omics Core and Genomics Platform for their help with RNASeq library sequencing and guidance on experimental design, the RI-INBRE/ SURF program for supporting summer undergraduate research for KCL, the URI Science and Engineering fellows program for supporting summer undergraduate research for KC and the Annual Mark Wilson conference attendees for many valuable suggestions and discussion.

## References

Anders, Simon, Paul Theodor Pyl, and Wolfgang Huber. 2015. “HTSeq--a Python Framework to Work with High-Throughput Sequencing Data.” Bioinformatics (Oxford, England) 31 (2): 166–69. https://doi.org/10.1093/bioinformatics/btu638.

Aziz, Ramy K., Daniela Bartels, Aaron A. Best, Matthew DeJongh, Terrence Disz, Robert A. Edwards, Kevin Formsma, et al. 2008. “The RAST Server: Rapid Annotations Using Subsystems Technology.” BMC Genomics 9 (February): 75. https://doi.org/10.1186/1471-2164-9-75.

Bernsen, J. 1986. “Dynamic Thresholding of Grey-Level Images.” Proc. 8th International Conf. on Pattern Recognition, Paris, France, Aug., 1986, 1251–55.

Bishai, W. R., N. S. Howard, J. A. Winkelstein, and H. O. Smith. 1994. “Characterization and Virulence Analysis of Catalase Mutants of Haemophilus Influenzae.” Infection and Immunity 62 (11): 4855–60. https://doi.org/10.1128/IAI.62.11.4855-4860.1994.

Brettin, Thomas, James J. Davis, Terry Disz, Robert A. Edwards, Svetlana Gerdes, Gary J. Olsen, Robert Olson, et al. 2015. “RASTtk: A Modular and Extensible Implementation of the RAST Algorithm for Building Custom Annotation Pipelines and Annotating Batches of Genomes.” Scientific Reports 5 (February): 8365. https://doi.org/10.1038/srep08365.

Chung, C. T., S. L. Niemela, and R. H. Miller. 1989. “One-Step Preparation of Competent Escherichia Coli: Transformation and Storage of Bacterial Cells in the Same Solution.” Proceedings of the National Academy of Sciences of the United States of America 86 (7): 2172–75. https://doi.org/10.1073/pnas.86.7.2172.

Cynamon, M. H., T. B. Sorg, and A. Patapow. 1988. “Utilization and Metabolism of NAD by Haemophilus Parainfluenzae.” Journal of General Microbiology 134 (10): 2789–99. https://doi.org/10.1099/00221287-134-10-2789.

Daims, Holger, Sebastian Lücker, and Michael Wagner. 2006. “Daime, a Novel Image Analysis Program for Microbial Ecology and Biofilm Research.” Environmental Microbiology 8 (2): 200–213. https://doi.org/10.1111/j.1462-2920.2005.00880.x.

Daims, Holger, and Michael Wagner. 2011. “In Situ Techniques and Digital Image Analysis Methods for Quantifying Spatial Localization Patterns of Nitrifiers and Other Microorganisms in Biofilm and Flocs.” Methods in Enzymology 496: 185–215. https://doi.org/10.1016/B978-0-12-386489-5.00008-7.

Deng, Yiqin, Chang Chen, Zhe Zhao, Jingjing Zhao, Annick Jacq, Xiaochun Huang, and Yiying Yang. 2016. “The RNA Chaperone Hfq Is Involved in Colony Morphology, Nutrient Utilization and Oxidative and Envelope Stress Response in Vibrio Alginolyticus.” PloS One 11 (9): e0163689. https://doi.org/10.1371/journal.pone.0163689.

Dewhirst, Floyd E., Tuste Chen, Jacques Izard, Bruce J. Paster, Anne C. R. Tanner, Wen-Han Yu, Abirami Lakshmanan, and William G. Wade. 2010. “The Human Oral Microbiome.” Journal of Bacteriology 192 (19): 5002–17. https://doi.org/10.1128/JB.00542-10.

Eren, A. Murat, Gary G. Borisy, Susan M. Huse, and Jessica L. Mark Welch. 2014. “Oligotyping Analysis of the Human Oral Microbiome.” Proceedings of the National Academy of Sciences of the United States of America 111 (28): E2875–2884. https://doi.org/10.1073/pnas.1409644111.

Eren, A. Murat, Hilary G. Morrison, Pamela J. Lescault, Julie Reveillaud, Joseph H. Vineis, and Mitchell L. Sogin. 2015. “Minimum Entropy Decomposition: Unsupervised Oligotyping for Sensitive Partitioning of High-Throughput Marker Gene Sequences.” The ISME Journal 9 (4): 968–79. https://doi.org/10.1038/ismej.2014.195.

Espinoza, Josh L., Derek M. Harkins, Manolito Torralba, Andres Gomez, Sarah K. Highlander, Marcus B. Jones, Pamela Leong, et al. 2018. “Supragingival Plaque Microbiome Ecology and Functional Potential in the Context of Health and Disease.” MBio 9 (6). https://doi.org/10.1128/mBio.01631-18.

Fantappiè, Laura, Matteo M. E. Metruccio, Kate L. Seib, Francesca Oriente, Elena Cartocci, Francesca Ferlicca, Marzia M. Giuliani, Vincenzo Scarlato, and Isabel Delany. 2009. “The RNA Chaperone Hfq Is Involved in Stress Response and Virulence in Neisseria Meningitidis and Is a Pleiotropic Regulator of Protein Expression.” Infection and Immunity 77 (5): 1842–53. https://doi.org/10.1128/IAI.01216-08.

Ferrières, Lionel, Gaëlle Hémery, Toan Nham, Anne-Marie Guérout, Didier Mazel, Christophe Beloin, and Jean-Marc Ghigo. 2010. “Silent Mischief: Bacteriophage Mu Insertions Contaminate Products of Escherichia Coli Random Mutagenesis Performed Using Suicidal Transposon Delivery Plasmids Mobilized by Broad-Host-Range RP4 Conjugative Machinery.” Journal of Bacteriology 192 (24): 6418–27. https://doi.org/10.1128/JB.00621-10.

Gustavsson, N., A. Diez, and T. Nyström. 2002. “The Universal Stress Protein Paralogues of Escherichia Coli Are Co-Ordinately Regulated and Co-Operate in the Defence against DNA Damage.” Molecular Microbiology 43 (1): 107–17. https://doi.org/10.1046/j.1365-2958.2002.02720.x.

Harrison, Alistair, Lauren O. Bakaletz, and Robert S. Munson. 2012. “Haemophilus Influenzae and Oxidative Stress.” Frontiers in Cellular and Infection Microbiology 2: 40. https://doi.org/10.3389/fcimb.2012.00040.

Hoeven, J. S. van der, A. I. Toorop, and R. H. Mikx. 1978. “Symbiotic Relationship of Veillonella Alcalescens and Streptococcus Mutans in Dental Plaque in Gnotobiotic Rats.” Caries Research 12 (3): 142–47. https://doi.org/10.1159/000260324.

Human Microbiome Project Consortium. 2012. “Structure, Function and Diversity of the Healthy Human Microbiome.” Nature 486 (7402): 207–14. https://doi.org/10.1038/nature11234.

Ishikawa, Takahiko, Yoshimitsu Mizunoe, Shun-ichiro Kawabata, Akemi Takade, Mine Harada, Sun Nyunt Wai, and Shin-ichi Yoshida. 2003. “The Iron-Binding Protein Dps Confers Hydrogen Peroxide Stress Resistance to Campylobacter Jejuni.” Journal of Bacteriology 185 (3): 1010–17. https://doi.org/10.1128/jb.185.3.1010-1017.2003.

Izawa, S., K. Maeda, T. Miki, J. Mano, Y. Inoue, and A. Kimura. 1998. “Importance of Glucose-6-Phosphate Dehydrogenase in the Adaptive Response to Hydrogen Peroxide in Saccharomyces Cerevisiae.” The Biochemical Journal 330 ( Pt 2) (March): 811–17. https://doi.org/10.1042/bj3300811.

Jensen, Anders, Christian F. P. Scholz, and Mogens Kilian. 2016. “Re-Evaluation of the Taxonomy of the Mitis Group of the Genus Streptococcus Based on Whole Genome Phylogenetic Analyses, and Proposed Reclassification of Streptococcus Dentisani as Streptococcus Oralis Subsp. Dentisani Comb. Nov., Streptococcus Tigurinus as Streptococcus Oralis Subsp. Tigurinus Comb. Nov., and Streptococcus Oligofermentans as a Later Synonym of Streptococcus Cristatus.” International Journal of Systematic and Evolutionary Microbiology 66 (11): 4803–20. https://doi.org/10.1099/ijsem.0.001433.

Jorth, Peter, Keith H. Turner, Pinar Gumus, Nejat Nizam, Nurcan Buduneli, and Marvin Whiteley. 2014. “Metatranscriptomics of the Human Oral Microbiome during Health and Disease.” MBio 5 (2): e01012–01014. https://doi.org/10.1128/mBio.01012-14.

Juneau, Richard A., Bing Pang, Chelsie E. Armbruster, Kyle A. Murrah, Antonia C. Perez, and W. Edward Swords. 2015. “Peroxiredoxin-Glutaredoxin and Catalase Promote Resistance of Nontypeable Haemophilus Influenzae 86-028NP to Oxidants and Survival within Neutrophil Extracellular Traps.” Infection and Immunity 83 (1): 239–46. https://doi.org/10.1128/IAI.02390-14.

Kamenšek, Simona, Zdravko Podlesek, Osnat Gillor, and Darja Zgur-Bertok. 2010. “Genes Regulated by the Escherichia Coli SOS Repressor LexA Exhibit Heterogeneous Expression.” BMC Microbiology 10 (November): 283. https://doi.org/10.1186/1471-2180-10-283.

Kanehisa, Minoru, Yoko Sato, and Kanae Morishima. 2016. “BlastKOALA and GhostKOALA: KEGG Tools for Functional Characterization of Genome and Metagenome Sequences.” Journal of Molecular Biology 428 (4): 726–31. https://doi.org/10.1016/j.jmb.2015.11.006.

Kapur, J.N., P.K. Sahoo, and A.K.C. Wong. 1985. “A New Method for Gray-Level Picture Thresholding Using the Entropy of the Histogram.” Computer Vision, Graphics, and Image Processing 29 (3): 273–85. https://doi.org/10.1016/0734-189X(85)90125-2.

Könönen, E., H. Jousimies-Somer, A. Bryk, T. Kilpi, and M. Kilian. 2002. “Establishment of Streptococci in the Upper Respiratory Tract: Longitudinal Changes in the Mouth and Nasopharynx up to 2 Years of Age.” Journal of Medical Microbiology 51 (9): 723–30. https://doi.org/10.1099/0022-1317-51-9-723.

Kosikowska, Urszula, Anna Biernasiuk, Paweł Rybojad, Renata Łoś, and Anna Malm. 2016. “Haemophilus Parainfluenzae as a Marker of the Upper Respiratory Tract Microbiota Changes under the Influence of Preoperative Prophylaxis with or without Postoperative Treatment in Patients with Lung Cancer.” BMC Microbiology 16 (April): 62. https://doi.org/10.1186/s12866-016-0679-6.

Kreth, Jens, Justin Merritt, and Fengxia Qi. 2009. “Bacterial and Host Interactions of Oral Streptococci.” DNA and Cell Biology 28 (8): 397–403. https://doi.org/10.1089/dna.2009.0868.

Langmead, Ben, and Steven L. Salzberg. 2012. “Fast Gapped-Read Alignment with Bowtie 2.” Nature Methods 9 (4): 357–59. https://doi.org/10.1038/nmeth.1923.

Liljemark, W. F., C. G. Bloomquist, L. A. Uhl, E. M. Schaffer, L. F. Wolff, B. L. Pihlstrom, and C. L. Bandt. 1984. “Distribution of Oral Haemophilus Species in Dental Plaque from a Large Adult Population.” Infection and Immunity 46 (3): 778–86. https://doi.org/10.1128/IAI.46.3.778-786.1984.

Liu, X., M. M. Ramsey, X. Chen, D. Koley, M. Whiteley, and A. J. Bard. 2011. “Real-Time Mapping of a Hydrogen Peroxide Concentration Profile across a Polymicrobial Bacterial Biofilm Using Scanning Electrochemical Microscopy.” Proceedings of the National Academy of Sciences 108 (7): 2668–73. https://doi.org/10.1073/pnas.1018391108.

Love, Michael I., Wolfgang Huber, and Simon Anders. 2014. “Moderated Estimation of Fold Change and Dispersion for RNA-Seq Data with DESeq2.” Genome Biology 15 (12): 550. https://doi.org/10.1186/s13059-014-0550-8.

Lundberg, B. E., R. E. Wolf, M. C. Dinauer, Y. Xu, and F. C. Fang. 1999. “Glucose 6-Phosphate Dehydrogenase Is Required for Salmonella Typhimurium Virulence and Resistance to Reactive Oxygen and Nitrogen Intermediates.” Infection and Immunity 67 (1): 436–38. https://doi.org/10.1128/IAI.67.1.436-438.1999.

Mark Welch, Jessica L., Floyd E. Dewhirst, and Gary G. Borisy. 2019. “Biogeography of the Oral Microbiome: The Site-Specialist Hypothesis.” Annual Review of Microbiology 73: 335–58. https://doi.org/10.1146/annurev-micro-090817-062503.

Mashimo, P. A., Y. Yamamoto, M. Nakamura, H. S. Reynolds, and R. J. Genco. 1985. “Lactic Acid Production by Oral Streptococcus Mitis Inhibits the Growth of Oral Capnocytophaga.” Journal of Periodontology 56 (9): 548–52. https://doi.org/10.1902/jop.1985.56.9.548.

Merritt, Judith H., Daniel E. Kadouri, and George A. O’Toole. 2005. “Growing and Analyzing Static Biofilms.” Current Protocols in Microbiology Chapter 1 (July): Unit 1B.1. https://doi.org/10.1002/9780471729259.mc01b01s00.

Mikx, F. H., and J. S. Van der Hoeven. 1975. “Symbiosis of Streptococcus Mutans and Veillonella Alcalescens in Mixed Continuous Cultures.” Archives of Oral Biology 20 (7): 407–10. https://doi.org/10.1016/0003-9969(75)90224-1.

Narayanan, Ajay M., Matthew M. Ramsey, Apollo Stacy, and Marvin Whiteley. 2017. “Defining Genetic Fitness Determinants and Creating Genomic Resources for an Oral Pathogen.” Applied and Environmental Microbiology 83 (14). https://doi.org/10.1128/AEM.00797-17.

Otsu, Nobuyuki. 1979. “A Threshold Selection Method from Gray-Level Histograms.” IEEE Transactions on Systems, Man, and Cybernetics 9 (1): 62–66. https://doi.org/10.1109/TSMC.1979.4310076.

Overbeek, Ross, Robert Olson, Gordon D. Pusch, Gary J. Olsen, James J. Davis, Terry Disz, Robert A. Edwards, et al. 2014. “The SEED and the Rapid Annotation of Microbial Genomes Using Subsystems Technology (RAST).” Nucleic Acids Research 42 (Database issue): D206–214. https://doi.org/10.1093/nar/gkt1226.

Pericone, C. D., K. Overweg, P. W. Hermans, and J. N. Weiser. 2000. “Inhibitory and Bactericidal Effects of Hydrogen Peroxide Production by Streptococcus Pneumoniae on Other Inhabitants of the Upper Respiratory Tract.” Infection and Immunity 68 (7): 3990–97. https://doi.org/10.1128/iai.68.7.3990-3997.2000.

Perkins, Arden, Kimberly J. Nelson, Derek Parsonage, Leslie B. Poole, and P. Andrew Karplus. 2015. “Peroxiredoxins: Guardians against Oxidative Stress and Modulators of Peroxide Signaling.” Trends in Biochemical Sciences 40 (8): 435–45. https://doi.org/10.1016/j.tibs.2015.05.001.

Ramsey, Matthew M., Marcelo O. Freire, Rebecca A. Gabrilska, Kendra P. Rumbaugh, and Katherine P. Lemon. 2016. “Staphylococcus Aureus Shifts toward Commensalism in Response to Corynebacterium Species.” Frontiers in Microbiology 7: 1230. https://doi.org/10.3389/fmicb.2016.01230.

Ramsey, Matthew M., Kendra P. Rumbaugh, and Marvin Whiteley. 2011. “Metabolite Cross-Feeding Enhances Virulence in a Model Polymicrobial Infection.” PLoS Pathogens 7 (3): e1002012. https://doi.org/10.1371/journal.ppat.1002012.

Redanz, Sylvio, Xingqun Cheng, Rodrigo A. Giacaman, Carmen S. Pfeifer, Justin Merritt, and Jens Kreth. 2018. “Live and Let Die: Hydrogen Peroxide Production by the Commensal Flora and Its Role in Maintaining a Symbiotic Microbiome.” Molecular Oral Microbiology 33 (5): 337–52. https://doi.org/10.1111/omi.12231.

Schäfer, A., A. Tauch, W. Jäger, J. Kalinowski, G. Thierbach, and A. Pühler. 1994. “Small Mobilizable Multi-Purpose Cloning Vectors Derived from the Escherichia Coli Plasmids PK18 and PK19: Selection of Defined Deletions in the Chromosome of Corynebacterium Glutamicum.” Gene 145 (1): 69–73. https://doi.org/10.1016/0378-1119(94)90324-7.

Schindelin, Johannes, Ignacio Arganda-Carreras, Erwin Frise, Verena Kaynig, Mark Longair, Tobias Pietzsch, Stephan Preibisch, et al. 2012. “Fiji: An Open-Source Platform for Biological-Image Analysis.” Nature Methods 9 (7): 676–82. https://doi.org/10.1038/nmeth.2019.

Segata, Nicola, Jacques Izard, Levi Waldron, Dirk Gevers, Larisa Miropolsky, Wendy S. Garrett, and Curtis Huttenhower. 2011. “Metagenomic Biomarker Discovery and Explanation.” Genome Biology 12 (6): R60. https://doi.org/10.1186/gb-2011-12-6-r60.

Segata, Nicola, Levi Waldron, Annalisa Ballarini, Vagheesh Narasimhan, Olivier Jousson, and Curtis Huttenhower. 2012. “Metagenomic Microbial Community Profiling Using Unique Clade-Specific Marker Genes.” Nature Methods 9 (8): 811–14. https://doi.org/10.1038/nmeth.2066.

Takashima, Eizo, and Kiyoshi Konishi. 2008. “Characterization of a Quinol Peroxidase Mutant in Aggregatibacter Actinomycetemcomitans.” FEMS Microbiology Letters 286 (1): 66–70. https://doi.org/10.1111/j.1574-6968.2008.01253.x.

Treerat, Puthayalai, Ulrike Redanz, Sylvio Redanz, Rodrigo A. Giacaman, Justin Merritt, and Jens Kreth. 2020. “Synergism between Corynebacterium and Streptococcus Sanguinis Reveals New Interactions between Oral Commensals.” The ISME Journal 14 (5): 1154–69. https://doi.org/10.1038/s41396-020-0598-2.

Wong, Sandy M. S., Kishore R. Alugupalli, Sanjay Ram, and Brian J. Akerley. 2007. “The ArcA Regulon and Oxidative Stress Resistance in Haemophilus Influenzae.” Molecular Microbiology 64 (5): 1375–90. https://doi.org/10.1111/j.1365-2958.2007.05747.x.

Xu, Yifan, Andreas Itzek, and Jens Kreth. 2014. “Comparison of Genes Required for H2O2 Resistance in Streptococcus Gordonii and Streptococcus Sanguinis.” Microbiology (Reading, England) 160 (Pt 12): 2627–38. https://doi.org/10.1099/mic.0.082156-0.

Zheng, Wenning, Tze King Tan, Ian C. Paterson, Naresh V. R. Mutha, Cheuk Chuen Siow, Shi Yang Tan, Lesley A. Old, Nicholas S. Jakubovics, and Siew Woh Choo. 2016. “StreptoBase: An Oral Streptococcus Mitis Group Genomic Resource and Analysis Platform.” PloS One 11 (5): e0151908. https://doi.org/10.1371/journal.pone.0151908.

Zhu, Bin, Lorna C. Macleod, Eric Newsome, Jinlin Liu, and Ping Xu. 2019. “Aggregatibacter Actinomycetemcomitans Mediates Protection of Porphyromonas Gingivalis from Streptococcus Sanguinis Hydrogen Peroxide Production in Multi-Species Biofilms.” Scientific Reports 9 (1): 4944. https://doi.org/10.1038/s41598-019-41467-9.

Zhu, Lin, and Jens Kreth. 2012. “The Role of Hydrogen Peroxide in Environmental Adaptation of Oral Microbial Communities.” Oxidative Medicine and Cellular Longevity 2012: 717843. https://doi.org/10.1155/2012/717843.

