## Supplemental Data for "Mechanisms underlying proximity between oral commensal bacteria"

Table S1: Minimum inhibitory concentrations in BHI-YE HP

| Strain | MIC (μM) |
|---|---|
| **Wildtype** | 834 |
| **Δ*katA*** | 834 |
| **Δ*ccp*** | 834 |
| **Δ*oxyR*** | 104 |

Table S2: *H. parainfluenzae* genes upregulated in *in vitro* coculture and *in vivo* metatranscriptome Espinoza et al., (2018)

| id | Description |
|---|---|
| fig\|6666666.571729.peg.60 | Glycerol uptake facilitator protein |
| fig\|6666666.571729.peg.32 | Type IV pilus biogenesis protein PilM |
| fig\|6666666.571729.peg.40 | Hydroxymethylpyrimidine ABC transporter transmembrane component |
| fig\|6666666.571729.peg.629 | NADP-specific glutamate dehydrogenase (EC 1.4.1.4);Ontology_term |
| fig\|6666666.571729.peg.1372 | ABC-type Fe3+-hydroxamate transport system periplasmic component |
| fig\|6666666.571729.peg.1681 | hypothetical protein |
| fig\|6666666.571729.peg.25 | AAA+ ATPase superfamily protein YifB/ComM associated with DNA recombination |
| fig\|6666666.571729.peg.687 | Phosphoribosylamine--glycine ligase (EC 6.3.4.13);Ontology_term |
| fig\|6666666.571729.peg.119 | Chloride channel protein EriC |
| fig\|6666666.571729.peg.95 | Phosphoribosylformylglycinamidine synthase synthetase subunit (EC 6.3.5.3) / Phosphoribosylformylglycinamidine synthase glutamine amidotransferase subunit (EC 6.3.5.3);Ontology_term |
| fig\|6666666.571729.peg.1446 | Allophanate hydrolase 2 subunit 2 (EC 3.5.1.54);Ontology_term |
| fig\|6666666.571729.peg.522 | 23S rRNA (uracil(1939)-C(5))-methyltransferase (EC 2.1.1.190);Ontology_term |
| fig\|6666666.571729.peg.173 | Putative ABC transporter of substrate X permease subunit I |
| fig\|6666666.571729.peg.686 | IMP cyclohydrolase (EC 3.5.4.10) / Phosphoribosylaminoimidazolecarboxamide formyltransferase (EC 2.1.2.3);Ontology_term |
| fig\|6666666.571729.peg.896 | 5'-deoxynucleotidase YfbR (EC 3.1.3.89);Ontology_term |
| fig\|6666666.571729.peg.242 | Fructose-16-bisphosphatase GlpX type (EC 3.1.3.11);Ontology_term |
| fig\|6666666.571729.peg.502 | hypothetical protein |
| fig\|6666666.571729.peg.866 | beta-galactosidase (EC 3.2.1.23);Ontology_term |
| fig\|6666666.571729.peg.3 | Methionine ABC transporter permease protein |
| fig\|6666666.571729.peg.792 | N-acetylmannosamine kinase (EC 2.7.1.60);Ontology_term |
| fig\|6666666.571729.peg.1078 | Aspartokinase (EC 2.7.2.4);Ontology_term |
| fig\|6666666.571729.peg.61 | Glycerol kinase (EC 2.7.1.30);Ontology_term |

Table S3: *H. parainfluenzae* genes downregulated in *in vitro* coculture and *in vivo* metatranscriptome Espinoza et al., (2018)

| id | Description |
|---|---|
| fig\|6666666.571729.peg.1245 | Molybdate-binding domain of ModE |
| fig\|6666666.571729.peg.903 | O-acetylhomoserine sulfhydrylase (EC 2.5.1.49) @ O-succinylhomoserine sulfhydrylase (EC 2.5.1.48);Ontology_term |
| fig\|6666666.571729.peg.1768 | hypothetical protein |
| fig\|6666666.571729.peg.905 | Chaperone protein DnaK |
| fig\|6666666.571729.peg.1409 | Outer membrane stress sensor protease DegQ serine protease |
| fig\|6666666.571729.peg.1340 | Transcription elongation factor GreA |
| fig\|6666666.571729.peg.1811 | Methionine repressor MetJ |
| fig\|6666666.571729.peg.1700 | Cytochrome c551 peroxidase (EC 1.11.1.5);Ontology_term |
| fig\|6666666.571729.peg.1850 | FIG002060: uncharacterized protein YggL |
| fig\|6666666.571729.peg.980 | hypothetical protein |
| fig\|6666666.571729.peg.430 | 33 kDa chaperonin HslO |
| fig\|6666666.571729.peg.904 | hypothetical protein |
| fig\|6666666.571729.peg.906 | hypothetical protein |
| fig\|6666666.571729.peg.665 | UPF0033 protein YeeD |
| fig\|6666666.571729.peg.1242 | Iron(III) ABC transporter solute-binding protein |
| fig\|6666666.571729.peg.1809 | Copper(I) chaperone CopZ |
| fig\|6666666.571729.peg.1012 | DedA protein |
| fig\|6666666.571729.peg.1818 | Heat shock protein 60 kDa family chaperone GroEL |
| fig\|6666666.571729.peg.67 | hypothetical protein |
| fig\|6666666.571729.peg.69 | hypothetical protein |
| fig\|6666666.571729.peg.496 | 2-haloalkanoic acid dehalogenase (EC 3.8.1.2);Ontology_term |
| fig\|6666666.571729.peg.57 | L-cystine ABC transporter substrate-binding protein TcyA |
| fig\|6666666.571729.peg.73 | Purine nucleoside phosphorylase (EC 2.4.2.1);Ontology_term |
| fig\|6666666.571729.peg.782 | LSU ribosomal protein L33p @ LSU ribosomal protein L33p zinc-independent |
| fig\|6666666.571729.peg.1316 | Stringent starvation protein B |
| fig\|6666666.571729.peg.1858 | Glutathionylspermidine synthase (EC 6.3.1.8) / Glutathionylspermidine amidohydrolase (EC 3.5.1.78);Ontology_term |
| fig\|6666666.571729.peg.46 | SSU ribosomal protein S21p |
| fig\|6666666.571729.peg.1797 | Soluble lytic murein transglycosylase (EC 4.2.2.n1) |
| fig\|6666666.571729.peg.1731 | Uncharacterized protein YbfG |
| fig\|6666666.571729.peg.1884 | Nucleoid-associated protein YaaK |
| fig\|6666666.571729.peg.717 | Uncharacterized protease YegQ |
| fig\|6666666.571729.peg.838 | Uracil permease @ Uracil:proton symporter UraA |
| fig\|6666666.571729.peg.1299 | LSU ribosomal protein L20p |

| | |
|---|---|
| fig\|6666666.571729.peg.1730 | Chaperone protein HscA |
| fig\|6666666.571729.peg.1861 | Phosphatidylserine decarboxylase (EC 4.1.1.65);Ontology_term |
| fig\|6666666.571729.peg.1800 | Glutamyl-tRNA synthetase (EC 6.1.1.17);Ontology_term |
| fig\|6666666.571729.peg.226 | Threonine dehydratase biosynthetic (EC 4.3.1.19);Ontology_term |
| fig\|6666666.571729.peg.1013 | LSU ribosomal protein L25p |
| fig\|6666666.571729.peg.948 | Ferric uptake regulation protein FUR |
| fig\|6666666.571729.peg.1305 | Threonyl-tRNA synthetase (EC 6.1.1.3);Ontology_term |

Table S4: *H. parainfluenzae* genes induced in *in vitro* coculture and *in vivo* metatranscriptome Jorth et al., (2014)

| id | Description |
|---|---|
| fig\|6666666.571729.peg.1083 | Peptide transport system ATP-binding protein sapF (TC 3.A.1.5.5) |
| fig\|6666666.571729.peg.747 | Type IV pilin PilA |
| fig\|6666666.571729.peg.656 | hypothetical protein |
| fig\|6666666.571729.peg.1288 | Type III restriction-modification system methylation subunit (EC 2.1.1.72);Ontology_term |
| fig\|6666666.571729.peg.40 | Hydroxymethylpyrimidine ABC transporter transmembrane component |
| fig\|6666666.571729.peg.1084 | Peptide ABC transporter ATP-binding protein SapD |
| fig\|6666666.571729.peg.1446 | Allophanate hydrolase 2 subunit 2 (EC 3.5.1.54);Ontology_term |
| fig\|6666666.571729.peg.25 | AAA+ ATPase superfamily protein YifB/ComM associated with DNA recombination |
| fig\|6666666.571729.peg.254 | hypothetical protein |
| fig\|6666666.571729.peg.1645 | DNA ligase |
| fig\|6666666.571729.peg.583 | FIGfam138462: Acyl-CoA synthetase AMP-(fatty) acid ligase |
| fig\|6666666.571729.peg.160 | hypothetical protein |
| fig\|6666666.571729.peg.1223 | Exodeoxyribonuclease V beta chain (EC 3.1.11.5);Ontology_term |
| fig\|6666666.571729.peg.629 | NADP-specific glutamate dehydrogenase (EC 1.4.1.4);Ontology_term |
| fig\|6666666.571729.peg.32 | Type IV pilus biogenesis protein PilM |
| fig\|6666666.571729.peg.60 | Glycerol uptake facilitator protein |
| fig\|6666666.571729.peg.1192 | Micrococcal nuclease (thermonuclease) homologs |
| fig\|6666666.571729.peg.95 | Phosphoribosylformylglycinamidine synthase synthetase subunit (EC 6.3.5.3) / Phosphoribosylformylglycinamidine synthase glutamine amidotransferase subunit (EC 6.3.5.3);Ontology_term |
| fig\|6666666.571729.peg.279 | Ribonuclease BN |
| fig\|6666666.571729.peg.687 | Phosphoribosylamine--glycine ligase (EC 6.3.4.13);Ontology_term |
| fig\|6666666.571729.peg.1141 | Succinyl-CoA ligase [ADP-forming] alpha chain (EC 6.2.1.5);Ontology_term |
| fig\|6666666.571729.peg.242 | Fructose-16-bisphosphatase GlpX type (EC 3.1.3.11);Ontology_term |
| fig\|6666666.571729.peg.102 | GALNS arylsulfatase regulator (Fe-S oxidoreductase) |
| fig\|6666666.571729.peg.382 | LSU rRNA pseudouridine(746) synthase (EC 5.4.99.29) @ tRNA pseudouridine(32) synthase (EC 5.4.99.28);Ontology_term |
| fig\|6666666.571729.peg.649 | hypothetical protein |
| fig\|6666666.571729.peg.1287 | Type III restriction-modification system restriction subunit (EC 3.1.21.5);Ontology_term |
| fig\|6666666.571729.peg.1163 | 14-alpha-glucan (glycogen) branching enzyme GH-13-type (EC 2.4.1.18);Ontology_term |
| fig\|6666666.571729.peg.1475 | Aerobic respiration control sensor protein ArcB (EC 2.7.13.3);Ontology_term |
| fig\|6666666.571729.peg.595 | FIG00847847: hypothetical protein |

| | |
|---|---|
| fig\|6666666.571729.peg.18 | Oligopeptide ABC transporter permease protein OppB (TC 3.A.1.5.1) |
| fig\|6666666.571729.peg.403 | hypothetical protein |
| fig\|6666666.571729.peg.61 | Glycerol kinase (EC 2.7.1.30);Ontology_term |
| fig\|6666666.571729.peg.592 | hypothetical protein |
| fig\|6666666.571729.peg.322 | Patatin-like phospholipase |
| fig\|6666666.571729.peg.593 | hypothetical protein |
| fig\|6666666.571729.peg.648 | Mobile element protein |
| fig\|6666666.571729.peg.597 | hypothetical protein |
| fig\|6666666.571729.peg.173 | Putative ABC transporter of substrate X permease subunit I |
| fig\|6666666.571729.peg.172 | Putative ABC transporter of substrate X ATP-binding subunit |
| fig\|6666666.571729.peg.1147 | UPF0115 protein YfcN |
| fig\|6666666.571729.peg.475 | Transcriptional regulator AsnC family |
| fig\|6666666.571729.peg.1258 | hypothetical protein |
| fig\|6666666.571729.peg.522 | 23S rRNA (uracil(1939)-C(5))-methyltransferase (EC 2.1.1.190);Ontology_term |
| fig\|6666666.571729.peg.119 | Chloride channel protein EriC |
| fig\|6666666.571729.peg.1122 | Phage integrase |
| fig\|6666666.571729.peg.1372 | ABC-type Fe3+-hydroxamate transport system periplasmic component |
| fig\|6666666.571729.peg.134 | Shikimate 5-dehydrogenase I alpha (EC 1.1.1.25);Ontology_term |
| fig\|6666666.571729.peg.1681 | hypothetical protein |
| fig\|6666666.571729.peg.252 | hypothetical protein |
| fig\|6666666.571729.peg.693 | hypothetical protein |
| fig\|6666666.571729.peg.249 | Lysophospholipase L2 (EC 3.1.1.5);Ontology_term |
| fig\|6666666.571729.peg.1636 | Putative DNA-binding protein in cluster with Type I restriction-modification system |
| fig\|6666666.571729.peg.501 | hypothetical protein |
| fig\|6666666.571729.peg.323 | DNA transformation protein TfoX |
| fig\|6666666.571729.peg.1477 | Uncharacterized protein MSMEG_6412 |
| fig\|6666666.571729.peg.969 | Trehalose operon transcriptional repressor |
| fig\|6666666.571729.peg.1916 | Glycosyl transferase |
| fig\|6666666.571729.peg.423 | deoxyribose-phosphate aldolase( EC:4.1.2.4 );Ontology_term |
| fig\|6666666.571729.peg.161 | hypothetical protein |
| fig\|6666666.571729.peg.1209 | tRNA pseudouridine(65) synthase (EC 5.4.99.26);Ontology_term |
| fig\|6666666.571729.peg.720 | hypothetical protein |
| fig\|6666666.571729.peg.730 | Rod shape-determining protein RodA |
| fig\|6666666.571729.peg.502 | hypothetical protein |
| fig\|6666666.571729.peg.855 | Glutamine synthetase adenylyl-L-tyrosine phosphorylase (EC 2.7.7.89) / Glutamate-ammonia-ligase adenylyltransferase (EC 2.7.7.42);Ontology_term |

| | |
|---|---|
| fig\|6666666.571729.peg.1078 | Aspartokinase (EC 2.7.2.4);Ontology_term |
| fig\|6666666.571729.peg.470 | hypothetical protein |
| fig\|6666666.571729.peg.525 | D-glycerate transporter (predicted) |
| fig\|6666666.571729.peg.1189 | Zn-ribbon-containing possibly nucleic-acid-binding protein |
| fig\|6666666.571729.peg.1786 | tRNA(Ile)-lysidine synthetase (EC 6.3.4.19);Ontology_term |
| fig\|6666666.571729.peg.866 | beta-galactosidase (EC 3.2.1.23);Ontology_term |
| fig\|6666666.571729.peg.960 | Inner membrane protein YedI |
| fig\|6666666.571729.peg.1017 | Cell division inhibitor Slr1223 (YfcH in EC) contains epimerase/dehydratase and DUF1731 domains |
| fig\|6666666.571729.peg.578 | FIG021862: membrane protein exporter |
| fig\|6666666.571729.peg.1011 | CRISPR-associated endonuclease Cas9 |
| fig\|6666666.571729.peg.176 | Sodium-dependent anion transporter family |
| fig\|6666666.571729.peg.1248 | Vitamin B12 ABC transporter permease protein BtuC |
| fig\|6666666.571729.peg.19 | Oligopeptide ABC transporter permease protein OppC (TC 3.A.1.5.1) |
| fig\|6666666.571729.peg.689 | hypothetical protein |
| fig\|6666666.571729.peg.1260 | hypothetical protein |
| fig\|6666666.571729.peg.258 | Tyrosine-specific transport protein |
| fig\|6666666.571729.peg.148 | Nitrate/nitrite response regulator protein NarP |
| fig\|6666666.571729.peg.21 | Oligopeptide ABC transporter ATP-binding protein OppF (TC 3.A.1.5.1) |
| fig\|6666666.571729.peg.1140 | Succinyl-CoA ligase [ADP-forming] beta chain (EC 6.2.1.5);Ontology_term |
| fig\|6666666.571729.peg.1235 | Oligopeptide/dipeptide ABC transporter permease protein / Oligopeptide/dipeptide ABC transporter ATP-binding protein |
| fig\|6666666.571729.peg.1631 | AAA ATPase central region |
| fig\|6666666.571729.peg.763 | Phosphatidate cytidylyltransferase (EC 2.7.7.41);Ontology_term |
| fig\|6666666.571729.peg.653 | Phosphoserine phosphatase (EC 3.1.3.3);Ontology_term |
| fig\|6666666.571729.peg.1236 | Oligopeptide ABC transporter ATP-binding protein OppF (TC 3.A.1.5.1) |
| fig\|6666666.571729.peg.104 | Choline-sulfatase (EC 3.1.6.6);Ontology_term |
| fig\|6666666.571729.peg.965 | Permease of the drug/metabolite transporter (DMT) superfamily |

Table S5: *H. parainfluenzae* genes repressed in *in vitro* coculture and *in vivo* metatranscriptome Jorth et al., (2014)

| id | Description |
|---|---|
| fig\|6666666.571729.peg.1646 | Thioredoxin |
| fig\|6666666.571729.peg.905 | Chaperone protein DnaK |
| fig\|6666666.571729.peg.1013 | LSU ribosomal protein L25p |
| fig\|6666666.571729.peg.1850 | FIG002060: uncharacterized protein YggL |
| fig\|6666666.571729.peg.1409 | Outer membrane stress sensor protease DegQ serine protease |
| fig\|6666666.571729.peg.1884 | Nucleoid-associated protein YaaK |
| fig\|6666666.571729.peg.73 | Purine nucleoside phosphorylase (EC 2.4.2.1);Ontology_term |
| fig\|6666666.571729.peg.948 | Ferric uptake regulation protein FUR |
| fig\|6666666.571729.peg.1793 | Membrane protein involved in the export of O-antigen and teichoic acid |
| fig\|6666666.571729.peg.903 | O-acetylhomoserine sulfhydrylase (EC 2.5.1.49) @ O-succinylhomoserine sulfhydrylase (EC 2.5.1.48);Ontology_term |
| fig\|6666666.571729.peg.338 | Hybrid peroxiredoxin hyPrx5 (EC 1.11.1.15);Ontology_term |
| fig\|6666666.571729.peg.1737 | tRNA (cytidine(32)/uridine(32)-2'-O)-methyltransferase (EC 2.1.1.200);Ontology_term |
| fig\|6666666.571729.peg.906 | hypothetical protein |
| fig\|6666666.571729.peg.1340 | Transcription elongation factor GreA |
| fig\|6666666.571729.peg.353 | hypothetical protein |
| fig\|6666666.571729.peg.1811 | Methionine repressor MetJ |
| fig\|6666666.571729.peg.1482 | hypothetical protein |
| fig\|6666666.571729.peg.1733 | Iron-sulfur cluster assembly iron binding protein IscA |
| fig\|6666666.571729.peg.1858 | Glutathionylspermidine synthase (EC 6.3.1.8) / Glutathionylspermidine amidohydrolase (EC 3.5.1.78);Ontology_term |
| fig\|6666666.571729.peg.142 | Ketol-acid reductoisomerase (NADP(+)) (EC 1.1.1.86);Ontology_term |
| fig\|6666666.571729.peg.1108 | hypothetical protein |
| fig\|6666666.571729.peg.1168 | 67-dimethyl-8-ribityllumazine synthase (EC 2.5.1.78);Ontology_term |
| fig\|6666666.571729.peg.764 | Alanine transaminase (EC 2.6.1.2);Ontology_term |
| fig\|6666666.571729.peg.57 | L-cystine ABC transporter substrate-binding protein TcyA |
| fig\|6666666.571729.peg.430 | 33 kDa chaperonin HslO |
| fig\|6666666.571729.peg.904 | hypothetical protein |
| fig\|6666666.571729.peg.1728 | Protein IscX believed to be involved in assembly of Fe-S clusters |
| fig\|6666666.571729.peg.1929 | SOS-response repressor and protease LexA (EC 3.4.21.88);Ontology_term |
| fig\|6666666.571729.peg.1316 | Stringent starvation protein B |
| fig\|6666666.571729.peg.666 | UPF0394 inner membrane protein YeeE |
| fig\|6666666.571729.peg.632 | Twin-arginine translocation protein TatB |

| | |
|---|---|
| fig\|6666666.571729.peg.1123 | Pyruvate kinase (EC 2.7.1.40);Ontology_term |
| fig\|6666666.571729.peg.1464 | GTP cyclohydrolase II (EC 3.5.4.25);Ontology_term |
| fig\|6666666.571729.peg.1214 | 3-hydroxyacyl-[acyl-carrier-protein] dehydratase FabA form (EC 4.2.1.59) @ Trans-2-decenoyl-[acyl-carrier-protein] isomerase (EC 5.3.3.14);Ontology_term |
| fig\|6666666.571729.peg.226 | Threonine dehydratase biosynthetic (EC 4.3.1.19);Ontology_term |
| fig\|6666666.571729.peg.841 | Nitric-oxide reductase (EC 1.7.99.7) quinol-dependent;Ontology_term |
| fig\|6666666.571729.peg.1861 | Phosphatidylserine decarboxylase (EC 4.1.1.65);Ontology_term |
| fig\|6666666.571729.peg.980 | hypothetical protein |
| fig\|6666666.571729.peg.1303 | Translation initiation factor 3 |
| fig\|6666666.571729.peg.871 | DNA polymerase III chi subunit (EC 2.7.7.7);Ontology_term |
| fig\|6666666.571729.peg.359 | SSU ribosomal protein S19p (S15e) |
| fig\|6666666.571729.peg.1643 | Arginine ABC transporter ATP-binding protein ArtP |
| fig\|6666666.571729.peg.1741 | Tol biopolymer transport system TolR protein |
| fig\|6666666.571729.peg.1338 | 23S rRNA (uridine(2552)-2'-O)-methyltransferase (EC 2.1.1.166);Ontology_term |
| fig\|6666666.571729.peg.1516 | Thiol peroxidase Tpx-type (EC 1.11.1.15);Ontology_term |
| fig\|6666666.571729.peg.1644 | D-sedoheptulose 7-phosphate isomerase (EC 5.3.1.28);Ontology_term |
| fig\|6666666.571729.peg.1587 | Ribonucleotide reductase of class III (anaerobic) large subunit (EC 1.17.4.2);Ontology_term |
| fig\|6666666.571729.peg.69 | hypothetical protein |
| fig\|6666666.571729.peg.1305 | Threonyl-tRNA synthetase (EC 6.1.1.3);Ontology_term |
| fig\|6666666.571729.peg.989 | Iron compound ABC transporter permease protein |
| fig\|6666666.571729.peg.1341 | D-alanyl-D-alanine carboxypeptidase (EC 3.4.16.4);Ontology_term |
| fig\|6666666.571729.peg.1096 | Thiol:disulfide interchange protein DsbC |
| fig\|6666666.571729.peg.949 | Flavodoxin 1 |
| fig\|6666666.571729.peg.1494 | Deoxycytidine triphosphate deaminase (EC 3.5.4.13);Ontology_term |
| fig\|6666666.571729.peg.358 | LSU ribosomal protein L2p (L8e) |
| fig\|6666666.571729.peg.1602 | FIG024746: hypothetical protein |
| fig\|6666666.571729.peg.1729 | Ferredoxin 2Fe-2S |
| fig\|6666666.571729.peg.496 | 2-haloalkanoic acid dehalogenase (EC 3.8.1.2);Ontology_term |
| fig\|6666666.571729.peg.1566 | UPF0307 protein YjgA |

Table S6: *H. parainfluenzae* genes induced in *in vitro* coculture and both *in vivo* metatranscriptomes

| id | Description |
|---|---|
| fig\|6666666.571729.peg.60 | Glycerol uptake facilitator protein |
| fig\|6666666.571729.peg.32 | Type IV pilus biogenesis protein PilM |
| fig\|6666666.571729.peg.40 | Hydroxymethylpyrimidine ABC transporter transmembrane component |
| fig\|6666666.571729.peg.629 | NADP-specific glutamate dehydrogenase (EC 1.4.1.4);Ontology_term |
| fig\|6666666.571729.peg.1372 | ABC-type Fe3+-hydroxamate transport system periplasmic component |
| fig\|6666666.571729.peg.1681 | hypothetical protein |
| fig\|6666666.571729.peg.25 | AAA+ ATPase superfamily protein YifB/ComM associated with DNA recombination |
| fig\|6666666.571729.peg.687 | Phosphoribosylamine--glycine ligase (EC 6.3.4.13);Ontology_term |
| fig\|6666666.571729.peg.119 | Chloride channel protein EriC |
| fig\|6666666.571729.peg.95 | Phosphoribosylformylglycinamidine synthase synthetase subunit (EC 6.3.5.3) / Phosphoribosylformylglycinamidine synthase glutamine amidotransferase subunit (EC 6.3.5.3);Ontology_term |
| fig\|6666666.571729.peg.1446 | Allophanate hydrolase 2 subunit 2 (EC 3.5.1.54);Ontology_term |
| fig\|6666666.571729.peg.522 | 23S rRNA (uracil(1939)-C(5))-methyltransferase (EC 2.1.1.190);Ontology_term |
| fig\|6666666.571729.peg.173 | Putative ABC transporter of substrate X permease subunit I |
| fig\|6666666.571729.peg.242 | Fructose-16-bisphosphatase GlpX type (EC 3.1.3.11);Ontology_term |
| fig\|6666666.571729.peg.502 | hypothetical protein |
| fig\|6666666.571729.peg.866 | beta-galactosidase (EC 3.2.1.23);Ontology_term |
| fig\|6666666.571729.peg.1078 | Aspartokinase (EC 2.7.2.4);Ontology_term |
| fig\|6666666.571729.peg.61 | Glycerol kinase (EC 2.7.1.30);Ontology_term |

Table S7: *H. parainfluenzae* genes repressed in *in vitro* coculture and both *in vivo* metatranscriptomes

| id | Description |
|---|---|
| fig\|6666666.571729.peg.903 | O-acetylhomoserine sulfhydrylase (EC 2.5.1.49) @ O-succinylhomoserine sulfhydrylase (EC 2.5.1.48);Ontology_term |
| fig\|6666666.571729.peg.905 | Chaperone protein DnaK |
| fig\|6666666.571729.peg.1409 | Outer membrane stress sensor protease DegQ serine protease |
| fig\|6666666.571729.peg.1340 | Transcription elongation factor GreA |
| fig\|6666666.571729.peg.1811 | Methionine repressor MetJ |
| fig\|6666666.571729.peg.1850 | FIG002060: uncharacterized protein YggL |
| fig\|6666666.571729.peg.980 | hypothetical protein |
| fig\|6666666.571729.peg.430 | 33 kDa chaperonin HslO |
| fig\|6666666.571729.peg.904 | hypothetical protein |
| fig\|6666666.571729.peg.906 | hypothetical protein |
| fig\|6666666.571729.peg.69 | hypothetical protein |
| fig\|6666666.571729.peg.496 | 2-haloalkanoic acid dehalogenase (EC 3.8.1.2);Ontology_term |
| fig\|6666666.571729.peg.57 | L-cystine ABC transporter substrate-binding protein TcyA |
| fig\|6666666.571729.peg.73 | Purine nucleoside phosphorylase (EC 2.4.2.1);Ontology_term |
| fig\|6666666.571729.peg.1316 | Stringent starvation protein B |
| fig\|6666666.571729.peg.1858 | Glutathionylspermidine synthase (EC 6.3.1.8) / Glutathionylspermidine amidohydrolase (EC 3.5.1.78);Ontology_term |
| fig\|6666666.571729.peg.1884 | Nucleoid-associated protein YaaK |
| fig\|6666666.571729.peg.1861 | Phosphatidylserine decarboxylase (EC 4.1.1.65);Ontology_term |
| fig\|6666666.571729.peg.226 | Threonine dehydratase biosynthetic (EC 4.3.1.19);Ontology_term |
| fig\|6666666.571729.peg.1013 | LSU ribosomal protein L25p |
| fig\|6666666.571729.peg.948 | Ferric uptake regulation protein FUR |
| fig\|6666666.571729.peg.1305 | Threonyl-tRNA synthethase (EC 6.1.1.3);Ontology_term |

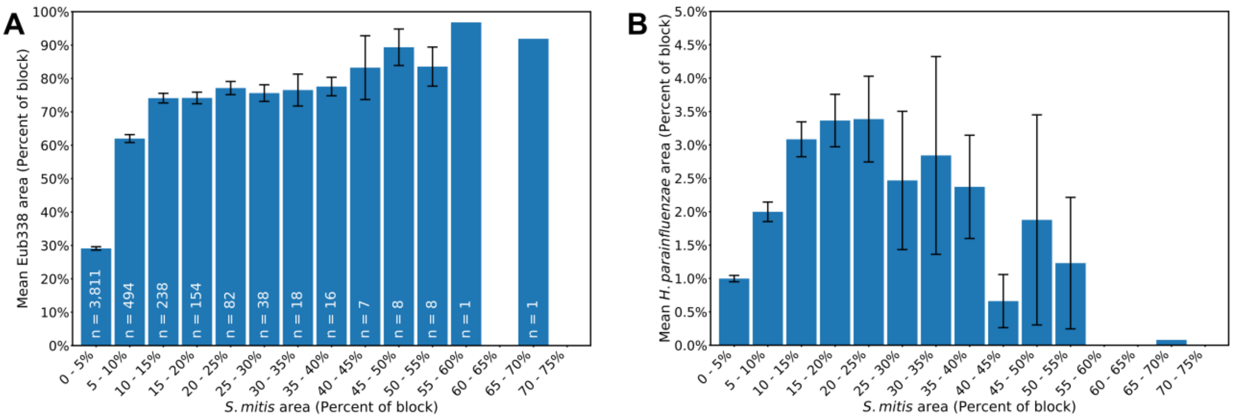


**Figure S1: *S. mitis* density relative to total bacteria and *H. parainfluenzae*** Mean local densities of (A) Eub338-labeled bacteria and (B) *H. parainfluenzae* with respect to the local density of *S. mitis* for 6.64 μm by 6.64 μm 4,876 blocks from 41 fields of view. The error bars represent ± 1 standard error.

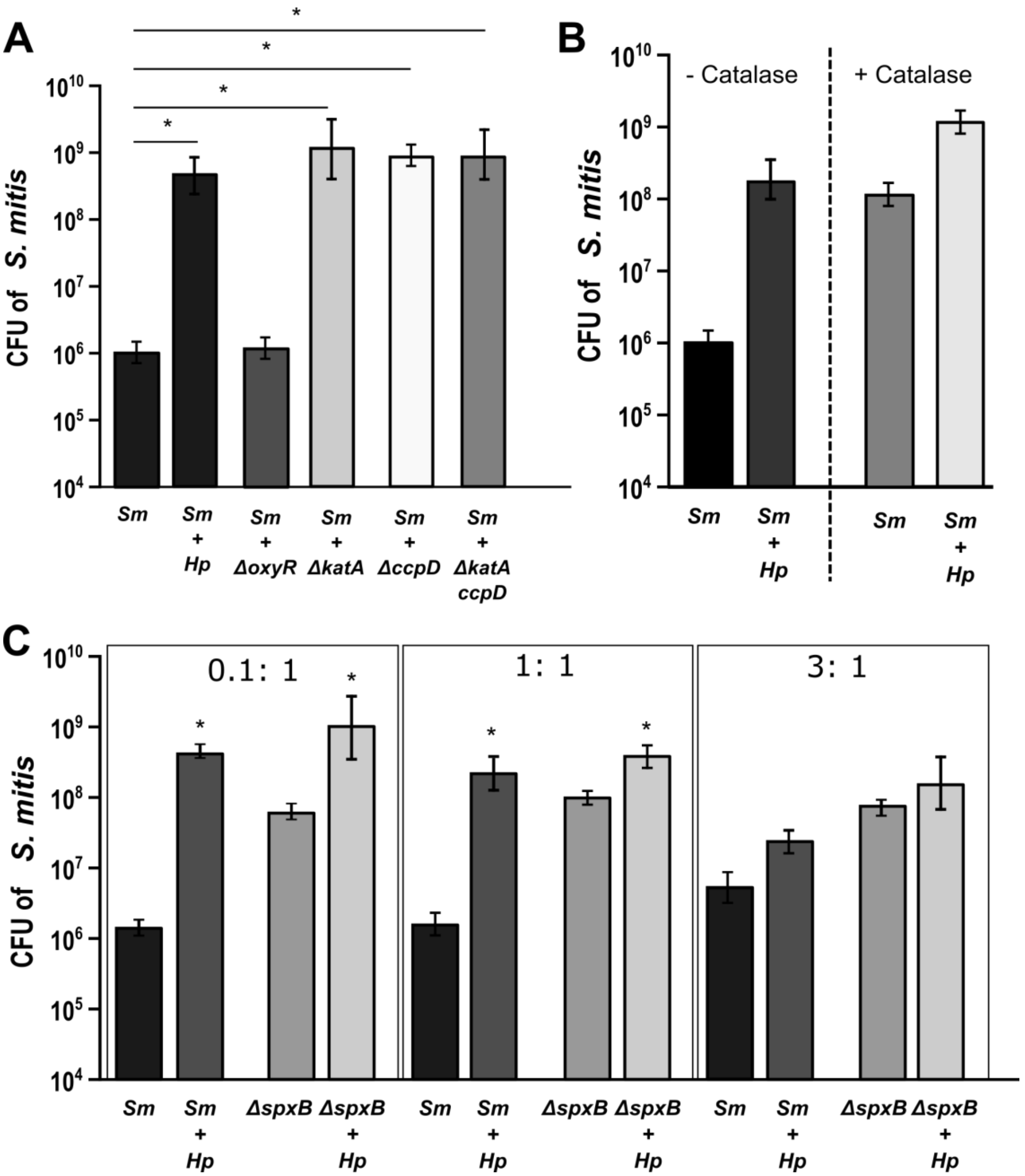


**Figure S2: *H. parainfluenzae* $H_2O_2$ detoxification aids *S. mitis* growth.** (A) *S. mitis* CFU when cocultured with *H. parainfluenzae* WT and indicated $H_2O_2$ resistance gene deletion mutants. Data are mean CFU, error bars indicate standard deviation for n=3. *denotes $p < 0.001$ by Student's t-test compared to monoculture CFU. (B) *S. mitis* CFU in mono and coculture with the addition of 20U/ml of exogenous catalase following incubation for 24 hours. (C) CFU counts of *S. mitis* (*Sm*) and the pyruvate oxidase mutant of *S. mitis* (*ΔspxB*) in mono and coculture with wildtype *H. parainfluenzae* (*Hp*). *Hp* had an initial inoculum of 4.65 x $10^6$ CFU/ml. Wildtype (*Sm*) and *S. mitis ΔspxB* with initial inoculums of 2.45x$10^5$, 1.55x$10^6$ or 3.45x$10^6$ CFU/ml. Data are mean CFU and error bars indicate standard deviation for n≥3. *denotes $p < 0.05$ by Student's t-test compared to monoculture.

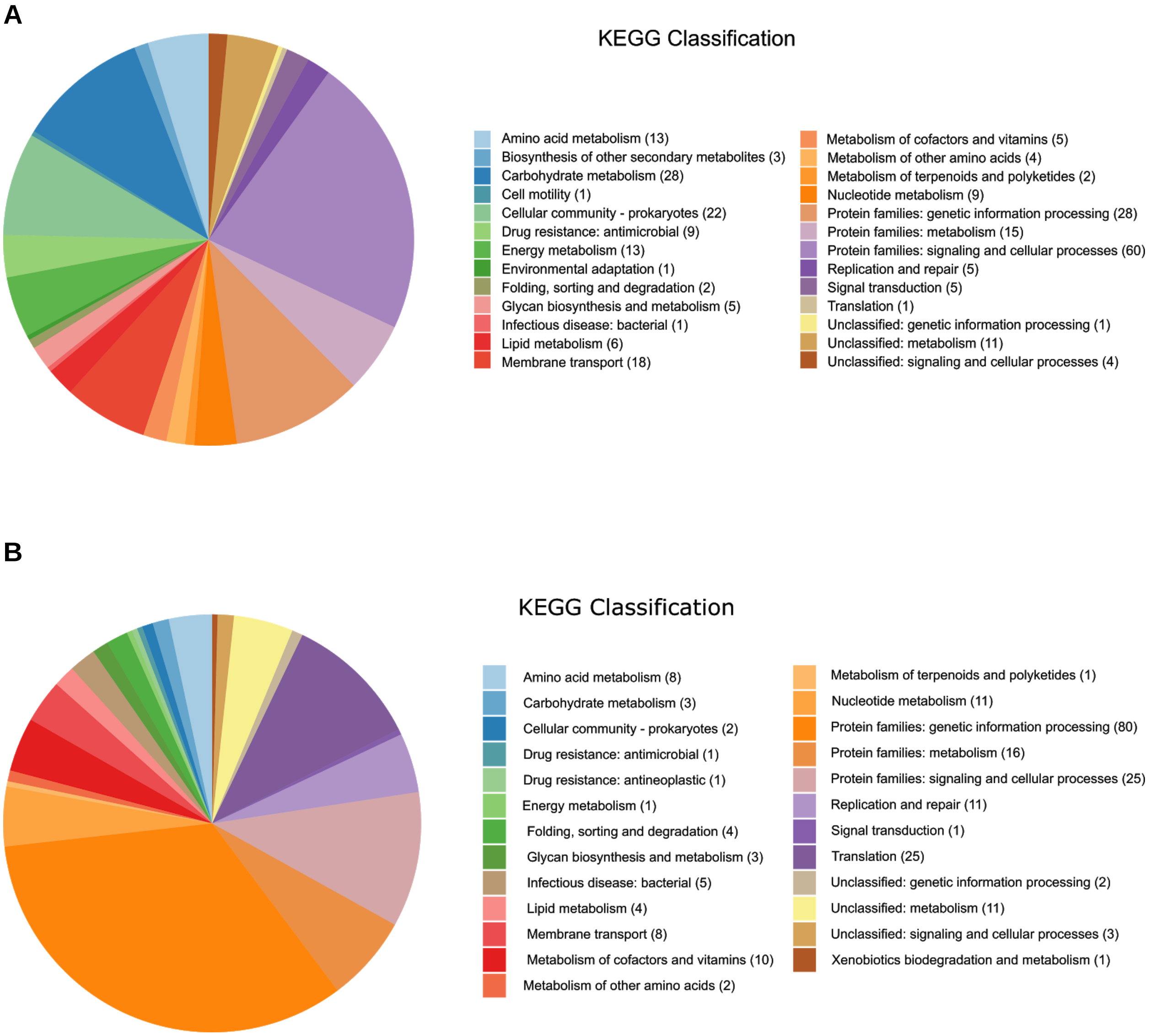


**Figure S3:** ***H. parainfluenzae*** **transcriptional response to** ***S. mitis*****. (A)** Pathways induced in *H. parainfluenzae* when cocultured with *S. mitis* based on number of genes that have > 2 fold increase in expression. **(B)** Pathways repressed when *H. parainfluenzae* is cocultured with *S. mitis* based on a 2 fold decrease in coculture.
